# Amino acid cross-feeding in *E. coli* globally and idiosyncratically alters mutant fitness

**DOI:** 10.64898/2026.05.27.727350

**Authors:** Duhita G. Sant, Tomas Lio Grudny, Atharv Jayprakash, Anastasia Dragan, Bidong D. Nguyen, Michael Manhart

**Affiliations:** Center for Advanced Biotechnology and Medicine, Rutgers University, Piscataway, NJ, USA; Institute of Integrative Biology, ETH Zurich, Zurich, Switzerland; Institute of Microbiology, ETH Zurich, Zurich, Switzerland; Department of Biochemistry and Molecular Biology, Robert Wood Johnson Medical School, Rutgers University, Piscataway, NJ, USA

**Keywords:** fitness, mutants, distribution of fitness effects, cross-feeding, amino acids, TnSeq, *E. coli*

## Abstract

Cross-feeding between microbes is believed to be common in many natural ecosystems, especially between species that are auxotrophic for essential nutrients such as amino acids. Several recent studies have demonstrated that cross-feeding and other ecological interactions between microbes alter their evolution and the fitness of mutations. However, these studies focused on interactions between different species, where the nature of the interaction is complex due to the large genetic differences between species, making it difficult to directly attribute changes in mutant fitness to a specific interaction. To address this problem, we use a synthetic cross-feeding interaction between isoleucine and methionine auxotrophs of *Escherichia coli*, two commonly observed auxotrophies in bacterial genomes. Using a library of genome-wide knockout mutants from transposon insertions, we measure the fitness of these mutants in the presence and absence of cross-feeding. We find that cross-feeding either isoleucine or methionine globally shifts mutant fitness to become more beneficial on average by inducing strong positive selection on a few mutants and rescuing strongly deleterious mutants. However, cross-feeding also affects mutants idiosyncratically: the most beneficial mutants under cross-feeding are neutral or deleterious without cross-feeding, and cross-feeding isoleucine affects different genes than cross-feeding methionine does. We discover one spontaneous mutant, with especially dramatic idiosyncratic effects under isoleucine cross-feeding, that achieves this phenotype by becoming a partial threonine auxotroph and reversing its ancestral isoleucine auxotrophy. This work directly demonstrates the statistical patterns and possible mechanisms by which a common ecological interaction between microbes alters their mutant fitness.

## INTRODUCTION

In diverse environments ranging from the human gut to environmental biofilms, microbes frequently exchange metabolic byproducts, known as cross-feeding [1–4]. A major driver of cross-feeding is when microbes are auxotrophic for essential nutrients, such as amino acids or vitamins, making secretions from other microbes in their environment a valuable source of these nutrients [2, 5]. Cross-feeding can be an important feature of an ecosystem’s state; for example, changes in cross-feeding within the human gut microbiome have been associated with gut health and several diseases [6–10].

This raises the question of how cross-feeding affects the fitness of mutations that arise in these ecosystems, and the consequences for evolution. A statistical approach to this question focuses on the distribution of fitness effects (DFE) across all spontaneous mutations in the genome [11], or a proxy for it. In some species, cross-feeding interactions tend to reduce the magnitude of mutant fitness effects [12–14]. A hypothesized mechanism is that deleterious mutants of the focal species are buffered by the partner species secreting compensating metabolites, while beneficial mutants are muted because growth of the focal species is always limited by its dependence on the partner species. This is equivalent to the putative weakest-link principle, which posits that adaptation of an interdependent community of species is limited by its least-adapted member [15, 16]. In other species, though, interactions have been found to change mutant fitness with no consistent bias [17– and some interactions appear not to significantly change mutant fitness at all [14, 21].

A challenge in interpreting these previous results, however, is that they all study interactions between different species, which often interact in multiple ways simultaneously (e.g., cross-feeding multiple nutrients and competing for others). The causality of these results from the underlying interactions is therefore difficult to determine, as it is unclear whether some interactions cause other interactions, and which interactions directly affect mutant fitness. An alternative approach, therefore, is to use strains of a single species with single-gene deletions that make them auxotrophs for essential nutrients [22– 26]. When two different such strains are grown together, they are obligate cross-feeders of the nutrients they are unable to synthesize, and since they are genetically identical besides single gene deletions causing the auxotrophy, they are limited in the number of other ways they can interact.

We use this approach to study how cross-feeding the amino acids isoleucine and methionine, two nutrients for which strains are commonly auxotrophic in nature [5, 6], affects mutant fitness in auxotrophic strains of *Escherichia coli* [26]. For each strain we generate libraries of random genome-wide transposon insertions that simulate spontaneous loss-of-function mutants [27, 28]. We find that cross-feeding isoleucine and cross-feeding methionine have similar and reproducible statistical effects on mutant fitness [29]: the average mutant fitness globally increases in both cases, with more strongly beneficial mutants and the most deleterious mutants being rescued by cross-feeding. Cross-feeding either amino acid also idiosyncratically transforms a few mutants that were neutral or deleterious in the absence of cross-feeding to become strongly beneficial in its presence. However, the specific genes affected by these idiosyncratic effects are different for cross-feeding isoleucine compared to crossfeeding methionine, indicating they are not generic consequences of cross-feeding or slow growth but specific effects of the auxotroph’s metabolic state. Finally, we investigate the mechanism of one mutant with especially strong idiosyncratic effects under isoleucine cross-feeding, which turns out to be caused by a phenotype that has a new partial auxotrophy for threonine and a partial reversion of the ancestral isoleucine auxotrophy.

## RESULTS

### A synthetic cross-feeding interaction to test statistical effects on the DFE

A recent theoretical study demonstrated that an ecological interaction can have two distinct types of statistical effects on the DFE [29]: global effects that transform fitness of all mutants the same way (e.g., shifting the mean fitness or rescaling the variance in fitness; Fig. 1A), and idiosyncratic effects that are unique to each mutant (Fig. 1B). This classification is useful because, besides being mathematically independent, the two types of effects correspond to distinct consequences for evolution. An interaction that changes the DFE globally alters evolution at the organismal scale, such as by changing the rate at which average organismal fitness increases. On the other hand, an interaction that has idiosyncratic effects on the DFE changes evolution at the molecular scale, by changing which genes and pathways adapt.

**FIG. 1.**
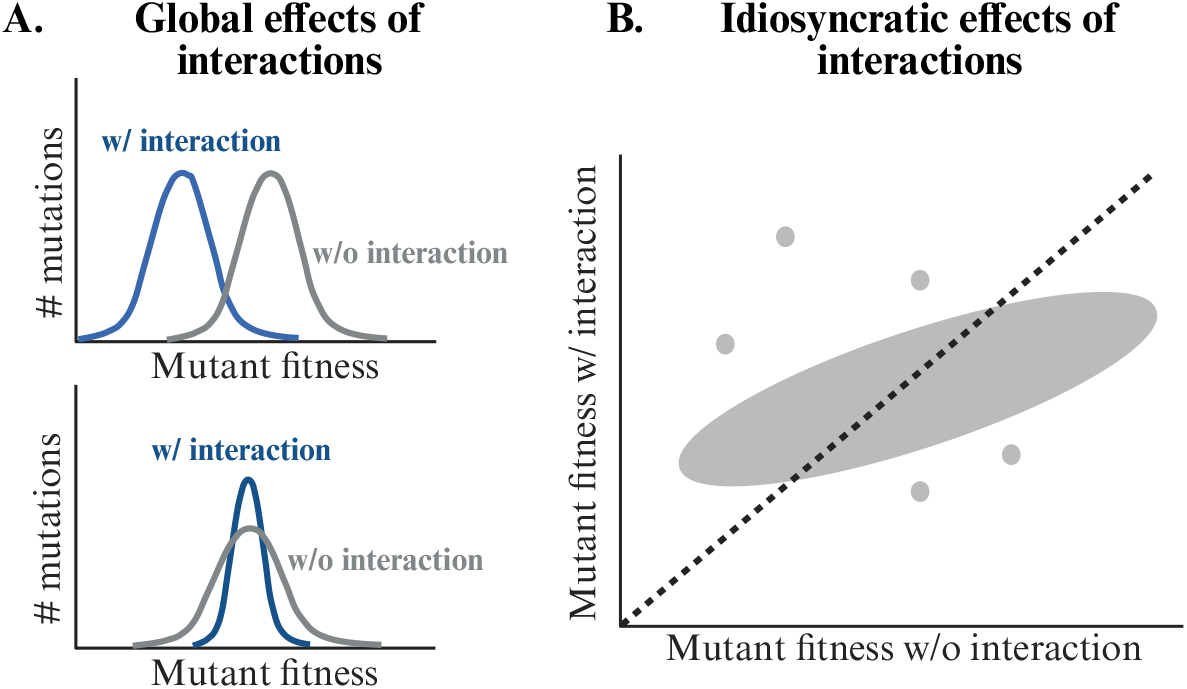
Potential statistical effects of cross-feeding interactions on the DFE. (A) Schematic DFEs with global effects of cross-feeding on the fitness mean (top panel) or fitness variance (bottom panel). (B) Schematic scatter plot of mutant fitness with and without a cross-feeding interaction, demonstrating idiosyncratic effects of the interaction on individual mutants.

To determine how a cross-feeding interaction generates these effects on the DFE, we focus on cross-feeding isoleucine and methionine, as previous work has predicted auxotrophs for these amino acids to be common in natural ecosystems [5, 6], and laboratory experiments have shown them to be reliably exchanged by auxotrophs [23, 26]. We use strains of *Escherichia coli* that previous work [26] engineered to be auxotrophs by deleting either the gene *ilvA* (for isoleucine) or *metA* (for methionine). As controls we use the prototrophic ancestors of each strain carrying the same fluorescent proteins (GFP for Δ*ilvA* and mCherry for Δ*metA*; Table S1). We first test that the auxotrophs stably coexist with each other in the absence of supplemented amino acids (confirming the presence of a mutualistic interaction) but that the prototrophs are neutral (Figs. S1 and S2, Methods). We then generate random-barcoded transposon insertion mutant libraries on the background of each of these four genotypes using established protocols [27, 28], so that each library contains many insertions per gene (Methods, Figs. 2, S3, and S4). Since insertion of the transposon within a gene typically disrupts the gene’s activity, these insertion mutants mimic the distribution of spontaneous loss-of-function mutations across the genome.

**FIG. 2.**
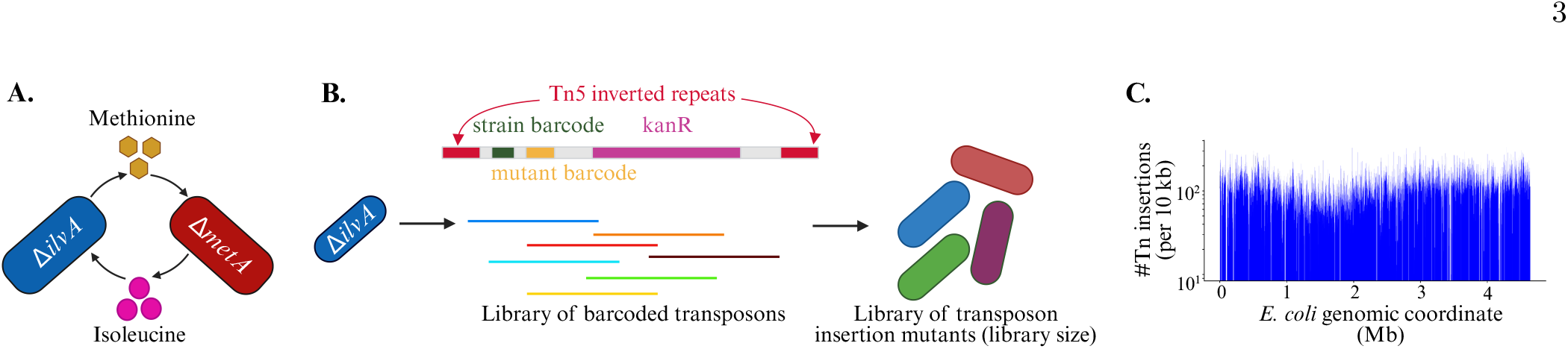
Model system for studying the effect of cross-feeding interactions on mutant fitness. (A) Synthetic cross-feeding interaction between isoleucine and methionine auxotrophs of *E. coli* [23, 26]. (B) Schematic of transposon sequence and workflow for generating random-barcoded transposon-insertion mutant libraries [27, 28] (on the background of Δ*ilvA* as an example; Methods). (C) Histogram of transposon insertion locations across the genome of Δ*ilvA* (see Fig. S3 for distributions for all libraries).

We perform mutant selection experiments in which each library is inoculated either with its ancestral genotype alone (“without interaction,” but with the necessary amino acid supplemented in the medium) or with both the ancestral genotype and the complementary auxotroph (“with interaction,” without any amino acids supplemented in the medium; Figs. 3 and S5, Table S2, Methods). To mimic the dynamics of a mutant arising spontaneously in a cross-feeding ecosystem, we inoculate the mutant library at low frequency relative to its ancestral genotype (so that the mutants must compete directly with their ancestral genotype) and with the cross-feeding auxotrophs at approximately the proportion at which they stably coexist (Fig. S1, Methods). We perform the mutant selection experiments over three serial transfers to more accurately infer each mutant’s dynamics. Over this time scale we find that the composition of the ancestral genotypes and the overall abundance of the mutant library as a whole do not significantly change (Fig. S6), indicating a stable ecological environment for the mutant selection. At each time point we sequence the mutants, determine the frequency, and infer fitness as the rate of change in their frequency (Figs. 3 and S5, Methods). With a few important exceptions we discuss below, replicate fitness measurements correlate well, indicating the robustness of this approach (Fig. S7 and Table S3).

**FIG. 3.**
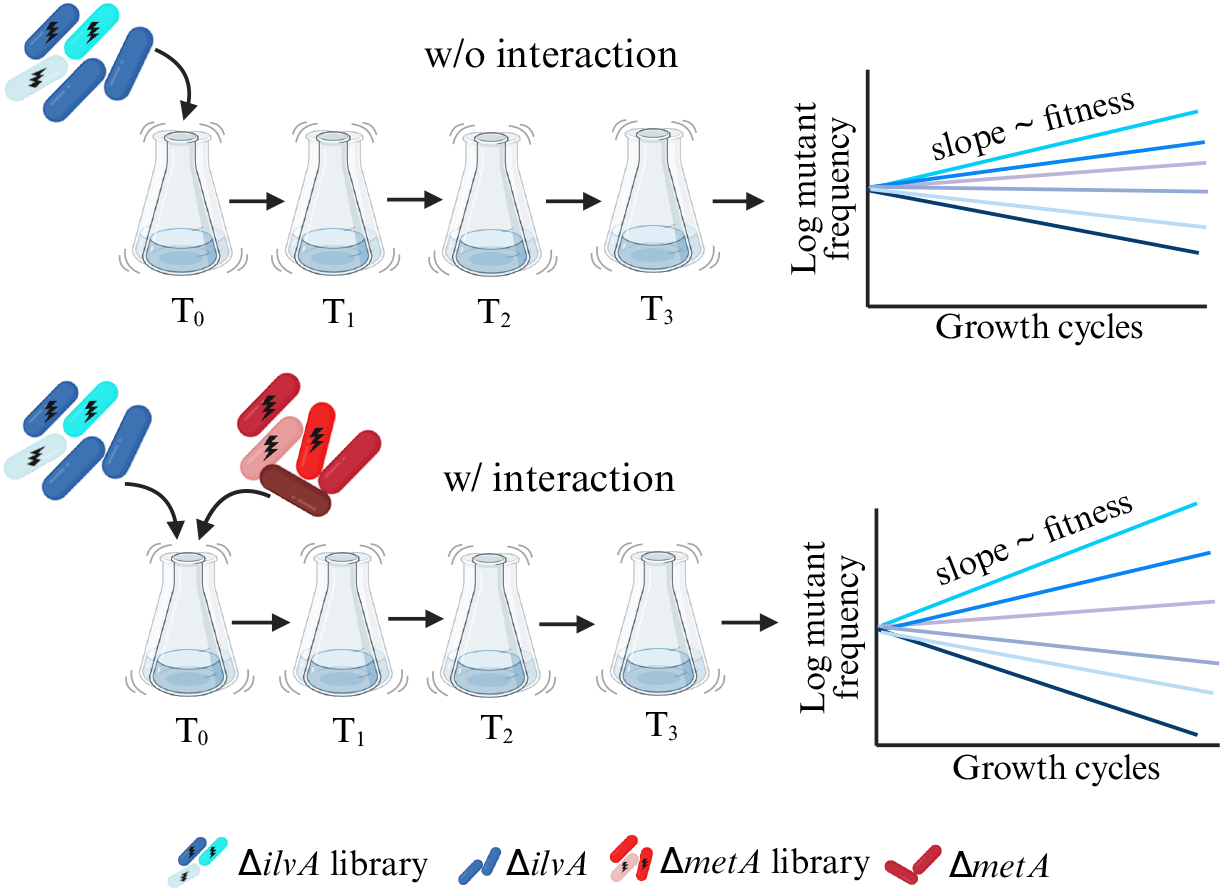
Schematic of mutant selection experiments with auxotrophs. Top row: selection experiment without the cross-feeding interaction, where the Δ*ilvA* mutant library is inoculated only with its ancestor Δ*ilvA* along with supplemented isoleucine, serially transferred over three batch growth cycles, and then sequenced at each time point to assemble mutant frequency trajectories (Methods). We also perform an analogous selection experiment with the Δ*metA* mutant library and its ancestor Δ*metA* without cross-feeding. Bottom row: same workflow but for selection experiment with the cross-feeding interaction, where the Δ*ilvA* mutant library is inoculated with both its ancestor Δ*ilvA* as well as Δ*metA*, without supplemented isoleucine. Figure S5 shows the analogous workflow for the prototroph controls.

### Cross-feeding isoleucine or methionine globally shifts the DFE to be more beneficial

Given fitness data across all mutants in the presence and absence of cross-feeding, we first consider cross-feeding’s global effects on the DFE as a whole (Fig. 4, Table S3). Crossfeeding globally shifts the DFE to eliminate the most deleterious effects and induce more strongly beneficial ones. There are also more weakly deleterious or nearly neutral mutants under cross-feeding. We can see these shifts reproducibly manifest in the mean and variance of the DFE (Fig. S8). These global effects on the DFE are common to cross-feeding either isoleucine (Fig. 4A) or methionine (Fig. 4B), suggesting they are a generic consequence of cross-feeding irrespective of its molecular details.

**FIG. 4.**
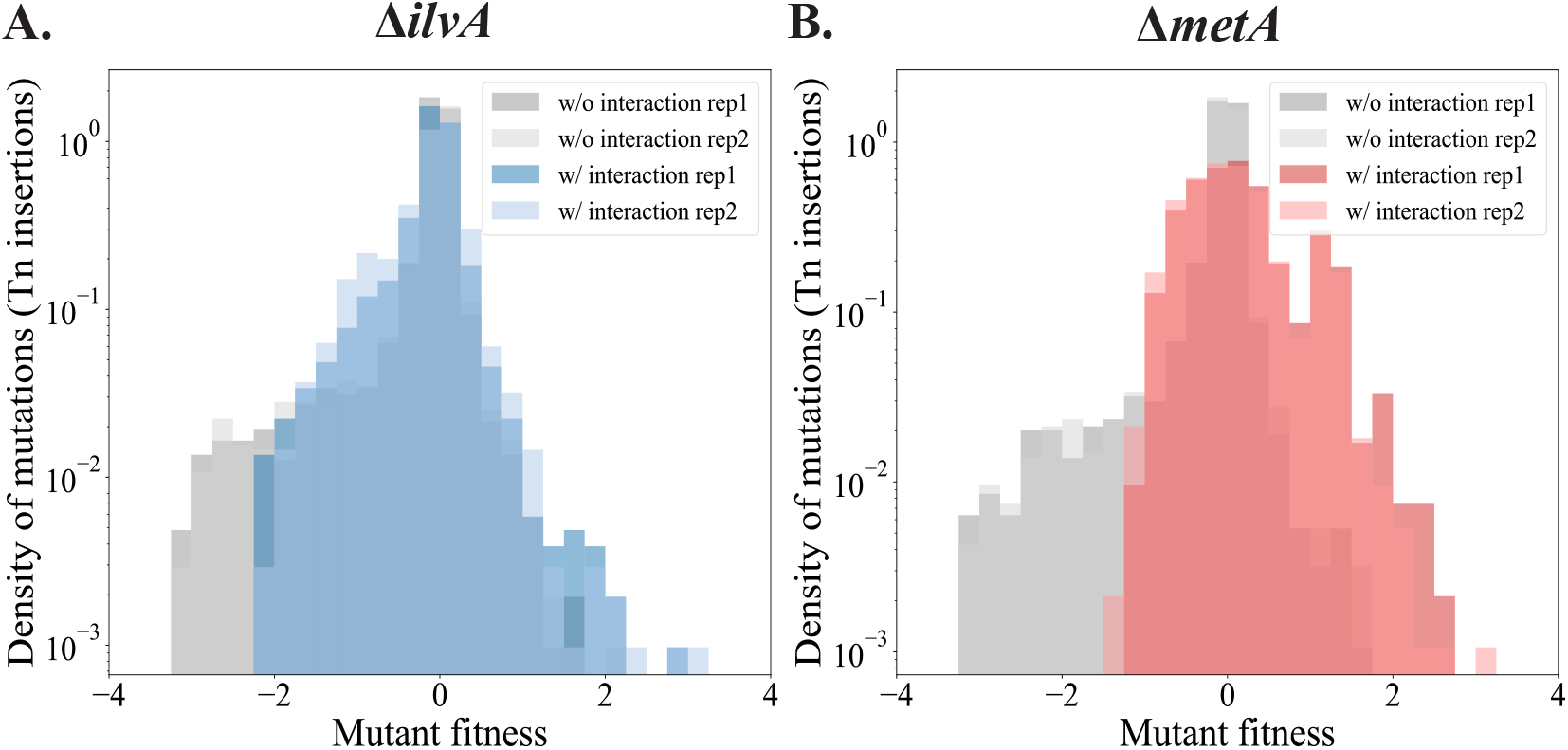
Cross-feeding either isoleucine or methionine globally shifts mutant fitness toward more beneficial effects. DFEs for the (A) Δ*ilvA* and (B) Δ*metA* mutant libraries, both with and without cross-feeding. Fitness is defined at the gene level (all insertions in the same gene are aggregated together; Methods). Figure S9 shows the analogous DFEs for the prototroph controls.

However, one caveat with this interpretation is that we observe a similar shift in the DFE in a control experiment in which we coculture the two prototrophic ancestors of Δ*ilvA* and Δ*metA* (having the same fluorescent markers but no gene deletions; Table S1). The expectation is that these prototrophs are phenotypically identical (differing only in fluorescent marker) and therefore their mutants will have the same fitness in monocultures as in a coculture with each other. For the GFP prototroph that serves as a control for Δ*ilvA*, we indeed see no significant shift in its DFE between a monoculture and a coculture with the mCherry prototroph (Fig. S9A). For the mCherry prototroph that is a control for Δ*metA*, however, we see a shift toward more beneficial mutants similar to the one observed for Δ*metA*, even though there is no crossfeeding interaction (compare Fig. 4B to Fig. S9B).

Why would mutants of the mCherry prototroph have different fitness in a coculture with a phenotypically identical strain? One major difference between the monoculture and coculture selection experiments is that we inoculated the mCherry prototroph at ∼ 20 times lower initial biomass in the coculture (Methods). We did this to match the setup of the cross-feeding coculture with the auxotrophs: in the mutant selection experiments with Δ*ilvA* and Δ*metA* cross-feeding, we inoculated those strains in the proportion 95:5 to mimic their composition at ecological equilibrium (Fig. S1). Thus, the mutants of Δ*ilvA* and its prototroph ancestor are at approximately the same biomass in both their monocultures and cocultures, whereas the mutants of Δ*metA* and its prototroph ancestor are diluted approximately 20 times in the coculture. This low abundance of the mutants in the cocultures may lead to undersampling of the most deleterious mutants, potentially explaining the apparent reduction in strongly deleterious effects for Δ*metA* in the cross-feeding coculture and the mCherry prototroph in its control coculture (Figs. 4B and S9B). However, that would not explain the appearance of strongly beneficial mutants for both Δ*metA* and the mCherry prototroph in their respective cocultures; instead, it raises the possibility that there are true frequency-dependent effects. A recent study of mutants from the Long-Term Evolution Experiment in *E. coli* found widespread frequencydependent selection on those mutants [30], with mutants generally having higher fitness at lower frequencies, consistent with our observations. As we will discuss in the next section, these strongly beneficial mutants in Δ*metA* are also in different genes than those in its prototroph ancestor, further suggesting that it is not merely an artifact of sampling low biomass.

### Cross-feeding idiosyncratically flips neutral or deleterious mutants to become strongly beneficial

The global effect of cross-feeding shifting the DFE toward higher fitness (Fig. 4) could occur by all mutants being equally shifted. However, we see that cross-feeding has idiosyncratic effects [29], with individual mutants being affected differently by cross-feeding (Fig. 5). On the deleterious end of the DFE, we see a systematic rescuing of many of the most deleterious mutants, but on the beneficial end, we see that individual mutants are flipped from being neutral or deleterious without cross-feeding to strongly beneficial with cross-feeding (red points in Fig. 5). Note that the prototroph control data shows for individual mutants of the GFP prototroph a strong correlation in monoculture and in coculture with the mCherry prototroph (Fig. S10A), as expected, while the mCherry prototroph mutants are only moderately correlated between culture conditions (Fig. S10B), potentially due to the aforementioned sampling bias and frequency-dependent effects.

**FIG. 5.**
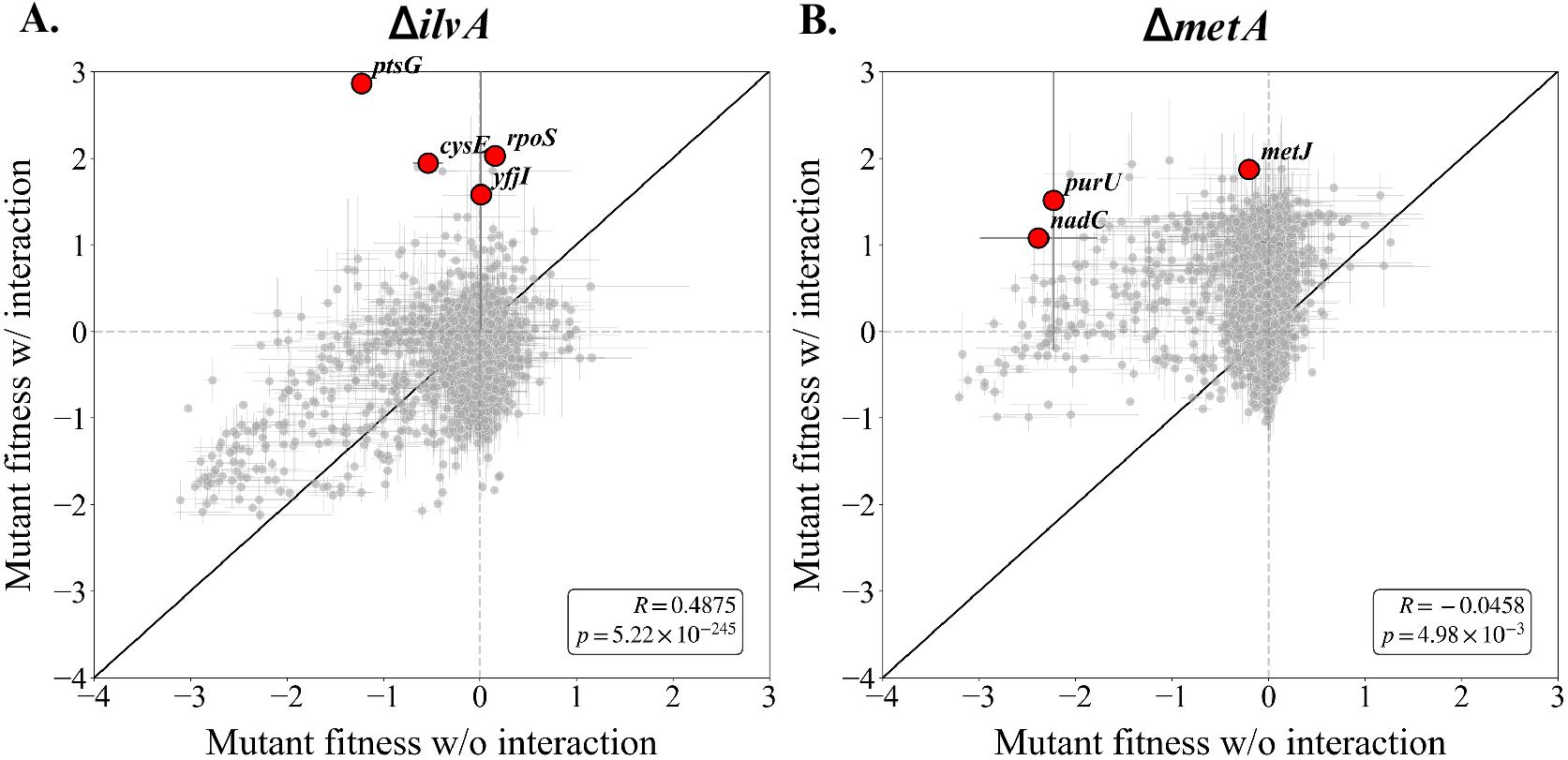
Cross-feeding either isoleucine or methionine idiosyncratically alters fitness of individual mutants. (A) Scatter plot of fitness for each gene (averaging all insertions in that gene; Methods) in Δ*ilvA* with and without isoleucine cross-feeding. (B) Same as (A) but for mutants in Δ*metA*. The legends show the Pearson correlation coefficients *R* and their associated *p*-values; the black diagonal line is the identity. Error bars represent standard error of the mean across two biological replicates (Fig. S7). Red points are mutants with strong beneficial effects under cross-feeding, with trajectories plotted in Fig. 6. Figure S10 shows the analogous scatter plots for the prototroph controls.

We can see these idiosyncratic effects more explicitly in the frequencies over time of these most beneficial mutants (Fig. 6; see Fig. S11 for the prototroph controls, and Fig. S12 for the same mutants across all other conditions). For example, insertions in the genes *ptsG* were neutral or deleterious on the background of Δ*ilvA* without cross-feeding (Fig. 6E,F), but became strongly beneficial when cross-feeding isoleucine (Fig. 6A,B). A previous study also found that transposon insertions in *ptsG*, encoding the permease of the glucose phosphotransferase transport system, were deleterious in monocultures but rescued in a cross-feeding culture [18] (with different cross-fed nutrients). While that study speculated that the change in selection on this gene was due to slower growth under cross-feeding, our reciprocal library approach shows that is not the case: if so, insertions in the same gene would be rescued on the Δ*metA* background as well (since both strains are presumably growing at the same rate in ecological equilibrium), but that is not the case. Figure S12 shows that insertions in *ptsG* are deleterious in Δ*metA*, cross-feeding or not, as well as on the prototroph backgrounds. Instead, the specific advantage of the *ptsG* insertion in Δ*ilvA* indicates it is an idiosyncratic effect of cross-feeding isolecuine.

**FIG. 6.**
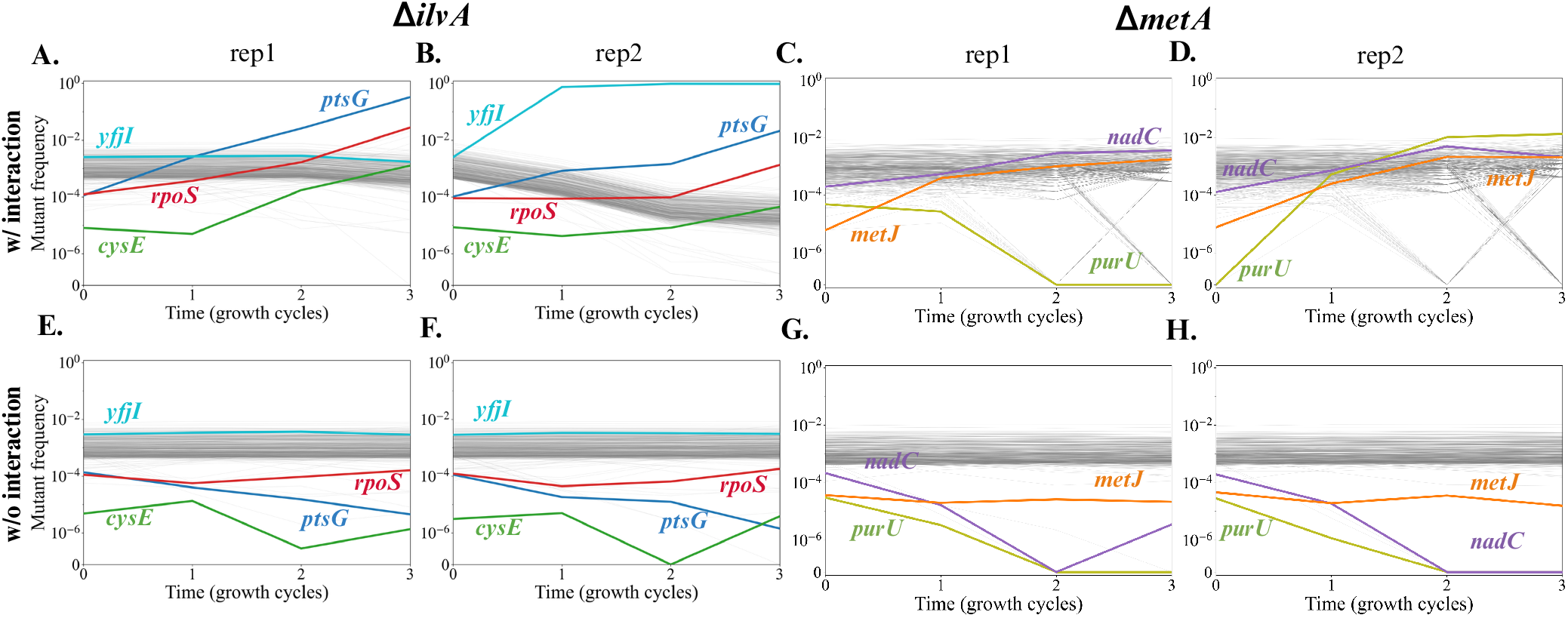
Dynamics of individual mutants with strong idiosyncratic effects from cross-feeding. Frequency over time for Δ*ilvA* (first and second columns) and Δ*metA* (third and fourth) mutants, with cross-feeding (first row) and without cross-feeding (second row). Colors and labels mark trajectories for the most beneficial mutants under cross-feeding, with grey trajectories being the remaining top 500 highest frequency mutants in each library. Figure S11 shows the analogous trajectories for the prototroph controls, and Fig. S12 shows all the same genes across conditions.

Two other genes with idiosyncratic fitness effects from isoleucine cross-feeding are *rpoS*, which mediates the general stress response, so that its inactivation may prevent costly, maladaptive stress behaviors in a stable coculture [18]; and *cysE*, in which insertions likely reduce L-cysteine levels, bypassing isoleucine auxotrophy by favoring the production of the 2-oxobutanoate precursor [31]. We observed a similar pattern of idiosyncratic effects in the Δ*metA* library but involving completely different genes (Fig. 6C,D,G,H; compare to Fig. S12). For methionine cross-feeding, insertions in *metJ* and *nadC* may have been deleterious in monocultures due to the loss of transcriptional regulation or redox capacity, they became highly beneficial while cross-feeding methionine, mirroring observations in other mutualistic systems where the presence of a partner leads to reduction in the selective pressure to maintain *de novo* biosynthesis of vitamins and precursors like NAD [14, 18].

### A putative mechanism for an idiosyncratic effect of isoleucine cross-feeding

A mutant on the background of Δ*ilvA* with especially dramatic idiosyncratic effects was an insertion in the gene *yfjI* (a prophage-associated protein of unknown function), which was consistently neutral without cross-feeding (Fig. 6E,F) but swept up to 74% after a single batch growth cycle and stabilized at 95–96% for the next two time points (Fig. 6B). However, this effect only occurred in a single replicate experiment (compare Fig. 6A and B) and in a single insertion (out of 789 unique insertions in *yfjI*) in *yfjI* (Fig. S13), suggesting that its strongly beneficial effect was driven not by the insertion itself but another genomic mutation linked to the insertion. We therefore isolated three colonies of this strain, which we verified by PCR of its barcode and Sanger sequencing (Methods). We performed whole-genome sequencing of these three isolates along with the Δ*ilvA* ancestor as a control (Methods), which revealed a nonsynonymous point mutation in *thrC* (threonine synthase) present in each *yfjI* insertion isolate but absent from the Δ*ilvA* ancestor (Table S4). We note that strong positive selection on an insertion in *purU* on the background of Δ*metA* accompanied the sweep of the *yfjI* insertion, suggesting either co-selection of both mutants together or a coincidental secondary mutation on the *purU* insertion strain as well.

The mutation in *thrC* suggested that its phenotype affected threonine metabolism. We therefore grew the three isolates of the *yfjI* insertion in minimal media with only isoleucine supplemented (to test the original auxotrophy), with only threonine (to test threonine auxotrophy), and with both isoleucine and methionine (Fig. 7A–C; see Fig. S14 for positive and negative controls). We found that the mutants grew poorly with only isoleucine but very well with only threonine compared to their Δ*ilvA* ancestor, suggested that the *thrC* mutation reversed the isoleucine auxotrophy but caused a new partial auxotrophy for threonine. The connection between metabolism for these amino acids is not surprising, since the *thrC* protein directly catalyzes the reaction producing threonine, which is the substrate for the *ilvA* protein [23] (and precursor to isoleucine further down the pathway).

**FIG. 7.**
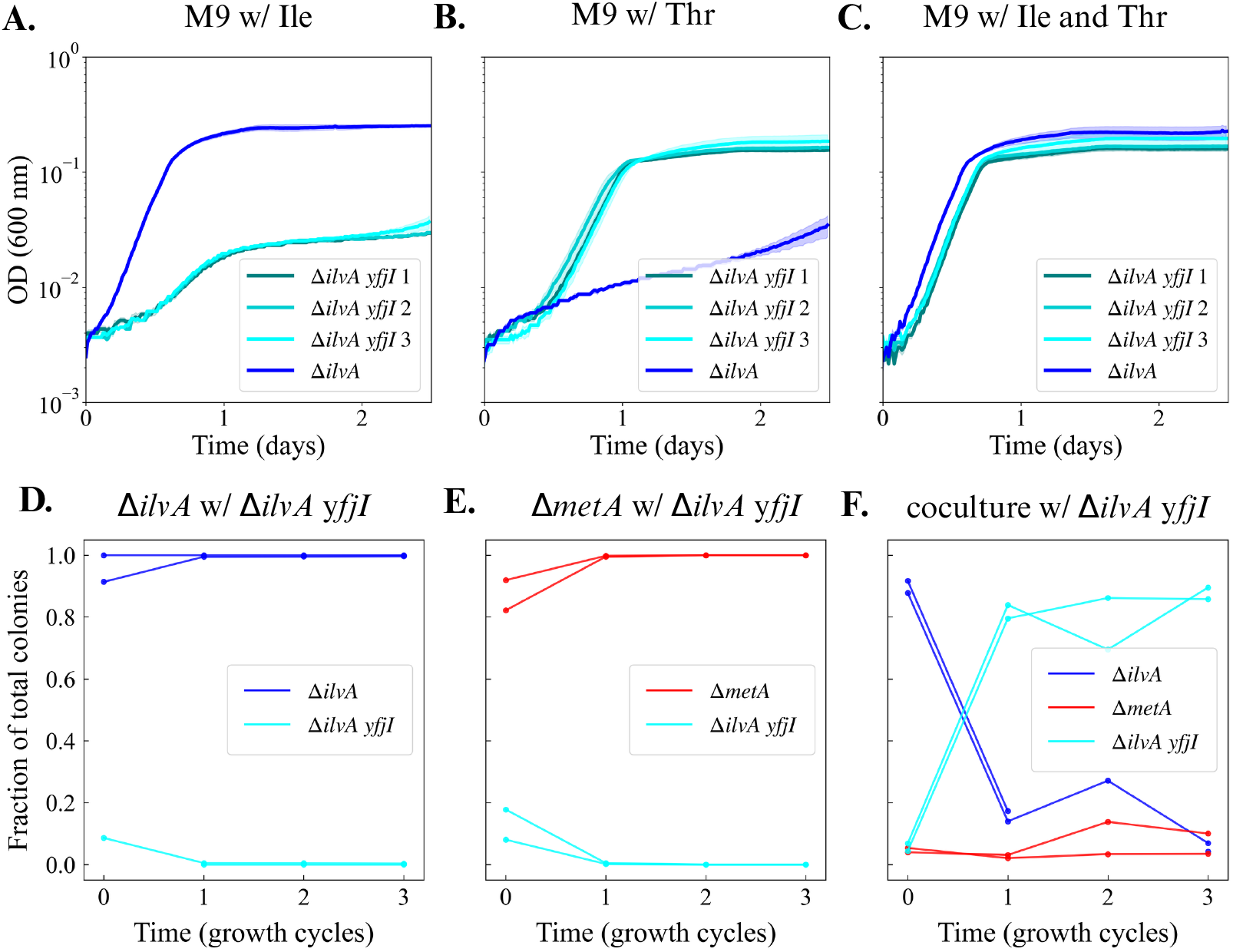
Secondary mutation in threonine synthase drives the idiosyncratic effect of isoleucine cross-feeding on the *yfjI* insertion strain. Growth curves of three isolates of the *yfjI* insertion strain and the Δ*ilvA* ancestor in minimal media supplemented with (A) isoleucine only, (B) threonine only, or (C) isoleucine and threonine (Methods). Dark line represents the mean and shading the standard error from three technical replicates (same inoculum). See Fig. S14 for positive and negative controls. Strain frequency trajectories from additional selection experiments with isolates of the *yfjI* insertion strain in (D) monoculture with Δ*ilvA* and supplemented isoleucine, (E) monoculture with Δ*metA* and supplemented isoleucine and methionine, and (F) coculture with Δ*ilvA* and Δ*metA* and no supplemented amino acids. Each trajectory has two biological replicates. Figure S15 shows the same data as estimated cell density (CFUs/mL).

We next directly tested that this mutant’s beneficial fitness is idiosyncratic to the cross-feeding coculture of Δ*ilvA* and Δ*metA*. Competing the mutant in monocultures with either Δ*ilvA* and Δ*metA* alone, the mutant was deleterious, dropping to low or zero frequency across replicates (Fig. 7D,E and Fig. S15A,B; see Fig. S16 for supporting spent media tests). However, in the coculture the mutant consistently rose to high abundance (Fig. 7F), but we observed that it could not totally outcompete either Δ*ilvA* and Δ*metA* (Fig. S15C), indicating that its fitness was frequency dependent. This is consistent with the observation from our original mutant selection experiments that this mutant did not entirely fix (maximum at 96% of all mutants, Fig. 6B), nor did it drive the mutant library as a whole to higher frequency relative to the Δ*ilvA* ancestor (Fig. S6).

## DISCUSSION

### Cross-feeding interactions globally shift the DFE towards beneficial fitness

This result advances the current understanding of the dynamic nature of the distribution of fitness effects (DFE) in response to the ecological context [11, 32]. On one hand, we observe a reduction of the deleterious tail, as described previously [14], that suggests metabolic rescue in which the presence of a partner provides essential metabolites that compensate for the metabolic deficiencies of a mutant [33, 34]. On the other hand, we also observe an enrichment of strongly beneficial mutants under cross-feeding. Altogether these results indicate that interactions can accelerate adaptation by creating more accessible evolutionary pathways [35]. This contrasts with the hypothesis that mutualistic interactions necessarily reduce the magnitude of fitness effects or constrain adaptation to the pace of the slowest community member [36].

### Change in the relative selection of mutants causes idiosyncratic effects in the presence of cross-feeding partner

While theoretical work has predicted that environment-specific selection can change the sign of a mutation’s fitness effect (from deleterious to beneficial or vice versa) [37, 38], empirical evidence in the context of metabolic cross-feeding has been sparse [15, 35]. Our identification of specific mutants that undergo a sign flip only in the presence of a partner provides a concrete molecular mechanism for these idiosyncratic effects.

The low correlation of mutant fitness with and without cross-feeding demonstrates that metabolic dependency is the primary driver of idiosyncratic effects on mutant fitness. In this context, mutations that would otherwise be purged by purifying selection are rescued or even promoted to high frequency because they complement the partner’s metabolic output. For example, the enrichment of *ptsG* knockouts in the Δ*ilvA* background suggests a specialized adaptation to the metabolic demands of acquiring isoleucine from a partner, rather than a general response to resource limitation, since we do not see positive selection on knockouts in the same gene on other genetic backgrounds. This implies that the partner creates a novel metabolic niche for this mutant. Such context-dependent selection highlights the idiosyncratic nature of metabolic niches created through cross-feeding.

### Metabolic rescue as a driver of diversity

The metabolic rescue observed in our study (the reduction of the deleterious tail) contributes to the growing literature on ecological buffering. While the Black Queen Hypothesis focuses on the adaptive loss of costly functions that are redundant due to public goods [39], our work suggests that partners also provide ecological buffering against the fitness costs of spontaneous, deleterious mutations. By reducing the deleterious tail of the DFE, cross-feeding allows a community to maintain higher genetic diversity, as fewer lineages are purged by purifying selection. A standout example of this rescue and subsequent adaptation is the strong idiosyncratic effect observed on the *thrC* mutation on the *yfjI* insertion strain. In isolation, the *thrC* mutant was rapidly purged when competing against ancestral strains without cross-feeding (Fig. 7D,E). However, in the presence of cross-feeding, this deleterious cost was transformed into a dominant selective advantage (Fig. 7F). The *thrC* mutation provided a fitness benefit by partially reversing the original isoleucine auxotrophy, a tradeoff that was only viable because the cross-feeding environment compensated for the mutant’s metabolic deficiencies. This reinforces the idea that metabolic rescue acts as a bridge, enabling populations to explore and fix complex, multi-step adaptive trajectories that would be purged by purifying selection in a purely competitive ecosystem.

### Evolution in cross-feeding communities

Our results suggest that evolution within a cross-feeding community is not merely an accelerated version of evolution in isolation, but a fundamentally distinct process altering both the tempo and the mode of evolution. The expansion of the beneficial side of the DFE implies that such systems will adapt more rapidly than isolated populations, as the partner provides a constant source of fuel for selection through niche construction. Furthermore, the observed metabolic rescue of deleterious mutations indicates that microbial interactions act as an evolutionary buffer, maintaining higher genetic diversity by preventing the immediate purging of lineages that would otherwise be lethal. This buffering allows the population to explore a more rugged fitness landscape, navigating evolutionary shortcuts where previously forbidden mutations, such as knockouts of *ptsG* or *nadC*, become viable assets. Ultimately, the lack of fitness correlation between treatments confirms that these systems do not evolve toward a universal optimum but toward an idiosyncratic, community-optimized state. This shift reinforces the paradigm that the presence of a metabolic partner unlocks new adaptive pathways, ensuring that the long-term evolutionary trajectory is tied to the stability and the output of the metabolic network [2, 35].

### Translation to natural systems

A critical question is how the dynamics observed in our engineered cross-feeding system translate to natural microbial communities. Although our model utilizes defined auxotrophs to enforce interaction, natural systems often feature more complex, spontaneously evolved dependencies. Extensive research has demonstrated that metabolic cross-feeding is a pervasive feature of microbial life, potentially arising through the Black Queen mechanism of adaptive gene loss [4, 40] In nature, the idiosyncratic fitness shifts we observed are likely amplified by the presence of multiple partners, and fluctuating nutrient availability. Our finding that cross-feeding interactions accelerate adaptation by reconfiguring the DFE aligns with empirical observations that metabolic dependencies drive rapid coevolution. Specifically, once a dependency is established, whether engineered or natural, it becomes selectively stabilized [41]. This relationship forces participants into a shared evolutionary fate where the fitness of one is tethered to the secretions of the other, mirroring the global and idiosyncratic effects seen in our auxotrophic libraries. However, a notable distinction exists regarding the robustness of these interactions. In our engineered system, the dependency is absolute; in natural systems, organisms may maintain degree of metabolic flexibility or engage in cheating behaviors. Yet, the high metabolic cost of de novo biosynthesis often makes reliance on a partner’s leaked metabolites more energetically favorable than self-sufficiency [41]. Our results provide the molecular resolution for this transition, showing that the beneficial shift in the DFE provides a powerful selective incentive for organisms to remain in states of dependency. Thus, our engineered model serves as a high-resolution lens through which we can view the same evolutionary forces that maintain stability and drive diversity in complex natural biomes [24].

## METHODS

### Strains

Table S1 lists the bacterial strains used in this study. Previous studies [25, 26] describe the construction of the auxotrophic *Escherichia coli* K-12 MG1655 strains Δ*ilvA*-GFP and Δ*metA*-mCherry and prototrophic *Escherichia coli* K-12 MG1655 TB204-GFP and TB205-mCherry.

### Preparation of barcoded transposon insertion libraries

We generated random barcoded transposon libraries in all four strains (Δ*ilvA* and Δ*metA* auxotrophs and TB204 and TB205 prototrophs) by electroporating transposomes as previously described [27, 28]. Breifly, we isolated a DNA fragment containing a kanamycin resistance gene (*aph/kanR*) from the pZ922 plasmid. We extracted the plasmid from *E. coli* DH5*α* using a commercial miniprep kit and cut out the resistance gene fragment via a NotI restriction digest. We purified the resulting 2.5 kb fragment by 1% (w/v) agarose gel electrophoresis to ensure the complete removal of supercoiled parental plasmid, which could otherwise contaminate downstream transformations. We generated functional transposons by PCR amplification of the purified kanR fragment, with primers designed to append the Tn5 inverted repeats (5^′^-CTGTCTCTTATACACATCT-3^′^), a library-specific 8-nt barcode, and a 20-nt random mutant barcode to the resistance gene. To ensure proper recognition by the transposase enzyme, both primers were synthesized with 5^′^ phosphate groups. We performed high-fidelity PCR using the Q5 Hot Start Master Mix supplemented with 4% DMSO to minimize secondary structures in the long primers. We purified the resulting ∼ 1.4 kb linear transposon products using a gel-based clean-up, taking care to minimize UV exposure and DNA cross-linking. We combined the purified transposon DNA with EZ-Tn5 transposase to form stable transposome complexes. We used a 2:1 stoichiometry of transposase to DNA and incubated the reaction at room temperature for 30 minutes.

We then prepared electrocompetent cells using a chilled washing protocol to achieve high transformation efficiency. Briefly, we grew *E. coli* cultures to midexponential phase (OD600 0.3—0.4) at 37°C. We then harvested the cells and subjected them to three successive washes in ice-cold 10% glycerol to remove residual salts and prevent arcing during electroporation. We performed all steps at 4°C with minimal mechanical stress to maximize cell viability. We generated the transposon-insertion mutant libraries by electroporating 50 *µ*L of electrocompetent cells with 1 *µ*L of the assembled transposome complexes. We performed electroporation at 1.8 kV using 1 mm cuvettes. Following a 2-hour recovery in SOC media at 37°C, we plated the cells onto LB agar supplemented with 0.05 g/L kanamycin. To ensure library quality, we used electroporation with the pZ922 plasmid as a positive control to verify transformation efficiency. We used electroporation of a transposase-negative reaction and screening of the main library on ampicillin to confirm the absence of contaminating parental plasmid as a negative control. We estimated the total library diversity by counting the number of unique colonies across many plates. The procedure yielded ∼ 5 × 10^4^–10^5^ unique transposon-insertion mutants. We harvested the colonies by flooding the plates with LB-glycerol, pooling them, and storing them in multiple aliquots at -70°C for downstream functional genomic assays.

### Sequencing and analysis of transposon insertion libraries

We processed raw Illumina paired-end sequencing reads (150 bp) to remove adapter sequences and low-quality bases using Trimmomatic in paired-end mode. We performed quality filtering using a sliding window approach (4 bp window), trimming reads where the average Phred quality score dropped below 15. We retained for downstream analysis only read pairs where both reads maintained a minimum length of 100 bp post-trimming. While trimming affected both reads, we observed and corrected a higher proportion of adapter contamination (∼ 15–18%) in the R2 reads. We screened the processed R1 reads for the Inverted Repeat (IR) and library barcodes. To account for potential sequencing errors or synthetic variations in barcode length, we identified the IR and library barcodes using flexible positional windows (bases 93—97 and 117–121, respectively). We extracted mutant barcodes from reads containing both a valid IR and library barcode in the expected orientations. To account for sequencing errors within the random mutant barcode sequences, we clustered the extracted barcodes using Bartender under default settings [42]. Following clustering, we appended the corrected barcode sequences to the FASTQ headers as record IDs. We discarded reads lacking the expected IR or library barcode sequences, maintaining paired-end synchrony by removing the corresponding R2 reads.

We aligned the processed reads, with IR and barcode sequences removed, to the *E. coli* reference genome using Bowtie2 in paired-end mode [43]. We employed an end-to-end alignment strategy to ensure high-stringency mapping of the genomic inserts. We converted the resulting SAM files to BAM format and coordinate-sorted and indexed them. We filtered the alignments to include only mapped, properly paired R1 reads. We categorized mapped reads by orientation (forward or reverse strand insertion) based on their mapping coordinates and reverse-complement status. We then extracted genomic coordinates and cross-referenced them with the *E. coli* reference genome to generate an initial insertion map. To resolve discrepancies where single barcodes mapped to multiple genomic locations (a known phenomenon in transposon sequencing), we assigned a barcode to a single definitive coordinate only if: 1) the coordinate contained at least 10 reads; 2) the primary coordinate accounted for ≥ 75% of the total reads for that barcode; and 3) the second-most frequent coordinate accounted for ≤ 12.5% of the total reads. For barcodes meeting these criteria, we consolidated all associated reads to the primary coordinate. To assess the randomness of the transposon library, we generated a null distribution by simulating random insertions across the genome. Finally, we tested the distribution of insertions across various genomic features. To investigate potential integration bias, we extracted 41-bp sequences centered on the insertion sites (± 20 bp) from the reference genome and analyzed them for GC-content and sequence motifs using sequence logos.

### Mutant selection experiments

We designed these experiments to measure mutant fitness by competing mutant libraries of the four strains with their ancestors in two distinct environments. The first environment is a monoculture without interaction, where mutants compete against their wild-type ancestors in M9 media supplemented with the necessary amino acids isoleucine or methionine. The second environment is a coculture with interaction, where mutants must compete against their ancestors in the presence of a different ancestral strain as well. We streaked frozen stocks of the ancestors on LB agar and grew them in M9 media to produce overnight cultures at an OD600 of at least 1. We resuspended frozen mutant libraries in 1X M9 salts to match the density of ancestral cultures. We washed all cultures three times in 1X M9 salts to remove residual media components before normalizing the cultures to a target OD600 of 1.

To inoculate the selection experiments, we combined normalized mutant libraries and ancestral wild-type strains in 500 mL flasks containing 148.5 mL of specific M9 minimal media (Table S2). We calibrated inoculation parameters to ensure that each unique mutant was represented by at least 10^2^ cells to minimize stochastic effects. For monocultures, we introduced a total inoculum volume of 1.5 mL at an OD600 of 1.0, maintaining a fixed initial mutant library frequency of 10% against 90% of the respective acenstral strain. In parallel, we inoculated the same volumes of cultures in microcentrifuge tubes, then pelleted and frozen them to serve as the initial time point for DNA extraction. We verified initial strain frequencies by serial dilution and plating on M9 agar (with and without kanamycin), and we then grew the cultures for 48 hours at 37°C with shaking. Every 48 hours, we transferred 1.5 mL of each culture into 148.5 mL of fresh media for a total of three growth cycles. At each transfer, we sampled the remaining culture for OD measurements, plated them to count colonies for strain frequency analysis (Fig. S6), and centrifuged them into pellets to be frozen for subsequent DNA sequencing.

### BarSeq sample preparation and sequencing

For BarSeq sample preparation, we extracted genomic DNA from frozen mutant library samples or enriched overnight cultures using the Qiagen DNeasy Blood and Tissue Kit. To ensure adequate mutant representation, we targeted approximately 3 × 10^9^ cells per sample for extraction, determined by measuring the OD600 of the cultures. Following purification, we measured DNA concentrations using both Nanodrop and Qubit instruments. We then normalized the concentration of purified DNA samples and submitted them for sequencing at Fasteris, with samples typically containing 40–80 ng/*µ*L of DNA in 25 *µ*L of nuclease-free water.

### Fitness data analysis and calculating DFEs

To process the BarSeq data, we multiplexed the raw sequencing reads to assign reads to specific experimental samples based on their multiplexing indices. We quantified the abundance of each mutant by identifying and counting the occurrences of unique 20-nt barcodes, which we subsequently aggregated by gene. To facilitate downstream statistical analysis and avoid undefined values during logarithmic transformation, a pseudocount of 0.1 was added to each barcode count. To estimate fitness of mutants relative to the ancestor without assuming the mean mutant fitness is zero, we normalized each barcode frequency relative to a set of putative pseudogenes that we expect to have zero fitness relative to the ancestor [44]. Specifically, we divided read counts for each gene in each sample by the average number of reads from fourteen reference genes: *gatR, insN, renD, yrhA, wbbL, ydbA, yqiG, yaiX, yhcE, insO, yaiT, nmpC, yejO*, and *yhiS*. We then transformed these normalized barcode frequencies by the logit function since this linearizes the trajectory of relative abundance over time under a null model of logistic dynamics [45]. To ensure statistical robustness, we calculated gene-level fitness as a weighted average based on barcode-level variance, with a maximum weight cap to prevent high-count barcodes from disproportionately biasing the results.

### Isolation of the *yfjI* insertion strain

We used replicate 2 of the auxotroph coculture mutant selection experiment in which the frequency of the *yfjI* insertion increased significantly (Fig. 6B), reaching 96% by the third time point. We streaked frozen stocks of this culture onto LB plates supplemented with 0.05 g/L kanamycin to obtain single colonies. Given the high prevalence of the *yfjI* insertion within the population, we anticipated that screening a limited number of individual colonies would yield the desired mutant with high probability. We performed the initial screening via colony PCR to identify candidates carrying the transposon insertion. We amplified the *yfjI* locus from three independent mutant replicates and the the ancestral Δ*ilvA* control. The thermal cycling protocol included an initial denaturation at 95°C for 2 minutes, followed by 30 cycles of denaturation (95°C for 30 seconds), annealing (56°C for 30 seconds), and extension (72°C for 2 minutes), with a final extension at 72°C for 5 minutes. We resolved the resulting amplicons on a 1% agarose gel and excised and purified the target bands using the QIAquick Gel Extraction Kit (Qiagen). We submitted the purified DNA samples to Genewiz for Sanger sequencing to confirm the insertion site and orientation. Finally, we extracted genomic DNA from the three validated *yfjI* colonies and the ancestral Δ*ilvA* control and submitted them for whole-genome sequencing (Genewiz) to find out any evidence of secondary mutations in *yfjI*.

### Growth experiments and pairwise competition assay with *yfjI* mutant

To evaluate the phenotypic consequences of the *yfjI* mutation, growth curve experiments were performed using the *yfjI* mutant and appropriate controls. Individual colonies were initially grown overnight in LB media supplemented with 0.05 g/L kanamycin. To remove residual media and nutrients, the overnight cultures were harvested and washed with 1X M9 salts. The cells were then normalized to an *OD*_600_ of 1.0 and inoculated into respective test media at an initial *OD*_600_ of 0.01. Based on sequencing results indicating a secondary mutation in the threonine synthase (*thrC*), the experimental media included M9 minimal media supplemented with threonine (M9 thr), isoleucine (M9 iso) or both (M9 iso + thr), with LB and M9 media serving as controls. Growth was monitored for 2.5 days using a LogPhase600 Reader (Agilent), with *OD*_600_ measurements recorded at 20-minute intervals under shaking at 37°C. The resulting raw data were processed using a custom Python script to perform blank subtraction and kinetic analysis, with the final growth curves visualized in Figure 7.

To investigate whether the *yfjI* mutant possesses a competitive advantage within a community context compared to monocultures, we characterized its growth dynamics in spent media derived from Δ*ilvA* and Δ*metA* monocultures and their corresponding coculture. Donor cultures were grown for 48 hours in M9 minimal media supplemented with either isoleucine (for Δ*ilvA*) or methionine (for Δ*metA*), while the coculture was inoculated at a 95:5 ratio (Δ*ilvA*:Δ*metA*) in unsupplemented M9 media similar to the Barseq experiment. Spent media was harvested via double centrifugation and sterilized using 0.2 *µ*m syringe filters. Three independent *yfjI* mutant replicates were grown overnight in LB supplemented with kanamycin, washed three times in 1× M9 salts to remove residual nutrients, and normalized to an *OD*_600_ of 1.0. The growth of these normalized *yfjI* cells was monitored for 2.5 days using a LogPhase600 Reader (Agilent). The resulting raw data were processed using a custom Python script to perform blank subtraction and kinetic analysis, with the final growth curves visualized in Figure 7.

To further validate the competitive advantage of the *yfjI* mutant, we performed direct pairwise and community competition assays over three growth cycles (six days). The *yfjI* mutant was competed against Δ*ilvA* and Δ*metA* in three distinct treatments: coculture (95% Δ*ilvA*, 5% Δ*metA*, with *yfjI* added at 10% of the Δ*ilvA* population) in M9 media without amino acids and two pairwise monoculture controls (90% auxotroph and 10% *yfjI* mutant) containing required amino acids. All strains were grown overnight, washed three times in 1× M9 salts to remove residual nutrients, and normalized to an *OD*_600_ of 1.0 before inoculation into M9 minimal media. The cultures were grown at 37°C with shaking and serially transferred every 48 hours (1.5 mL into 148.5 mL fresh media). To track the population dynamics, samples were taken at each transfer, serially diluted, and plated onto selective media: LB agar supplemented with kanamycin was used to quantify *yfjI* mutants, while M9 minimal agar supplemented with specific amino acids allowed for the enumeration of the total auxotroph populations. This longitudinal competition assay provided a direct quantitative measure of the fitness of *yfjI* mutant relative to Δ*ilvA* and Δ*metA* in monoculture and coculture environments.

## ACKNOWLEDGMENTS

This work was supported by the Swiss NSF (award PZ00P3 180147), NIGMS (award R35GM16022), and the Human Frontier Science Program (award RGEC30/2024).

## References

[1] M. A. Fischbach and J. L. Sonnenburg. Eating for two: How metabolism establishes interspecies interactions in the gut. Cell Host Microbe, 10:336–347, 2011.

[2] G. D’Souza, S. Shitut, D. Preussger, G. Yousif, S. Waschina, and C. Kost. Ecology and evolution of metabolic cross-feeding interactions in bacteria. Nat Prod Rep, 35:455–488, 2018.

[3] A. Goyal, T. Wang, V. Dubinkina, and S. Maslov. Ecology-guided prediction of cross-feeding interactions in the human gut microbiome. Nat Commun, 12:1335, 2021.

[4] C. Kost, K. R. Patil, J. Friedman, S. L. Garcia, and M. Ralser. Metabolic exchanges are ubiquitous in natural microbial communities. Nat Microbiol, 8:2244–2252, 2023.

[5] J. Ramoneda, T. B. N. Jensen, M. N. Price, E. O. Casamayor, and N. Fierer. Taxonomic and environmental distribution of bacterial amino acid auxotrophies. Nat Commun, 14:7608, 2023.

[6] S. Starke, D. M. M. Harris, J. Zimmermann, S. Schuchardt, M. Oumari, D. Frank, C. Bang, P. Rosenstiel, S. Schreiber, N. Frey, A. Franke, K. Aden, and S. Waschina. Amino acid auxotrophies in human gut bacteria are linked to higher microbiome diversity and long-term stability. ISME J, 17:2370–2380, 2023.

[7] V. R. Marcelino, C. Welsh, C. Diener, E. L. Gulliver, E. L. Rutten, R. B. Young, E. M. Giles, S. M. Gibbons, C. Greening, and S. C. Forster. Disease-specific loss of microbial cross-feeding interactions in the human gut. Nat Commun, 14:6546, 2023.

[8] I. Veseli, Y. T. Chen, M. S. Schechter, C. Vanni, E. C. Fogarty, A. R. Watson, B. Jabri, R. Blekhman, A. D. Willis, M. K. Yu, A. Fernandez-Guerra, J. Füssel, and A. M. Eren. Microbes with higher metabolic independence are enriched in human gut microbiomes under stress. bioRxiv, preprint, 2023-05-26. doi: 10.1101/2023.05.10.540289.

[9] A. R. Watson, J. Füssel, I. Veseli J. Z. DeLongchamp, M. Silva, F. Trigodet, K. Lolans, A. Shaiber, E. Fogarty, J. M. Runde, C. Quince, M. K. Yu, A. Söylev, H. G. Morrison, S. T. M. Lee, D. Kao, D. T. Rubin, B. Jabri, T. Louie, and A. M. Eren. Metabolic independence drives gut microbial colonization and resilience in health and disease. Genome Biol, 24:78, 2023.

[10] R. Corral López, J. A. Bonachela, M. G. Dominguez-Bello, M. Manhart, S. A. Levin, M. J. Blaser, and M. A. Muñoz. Imbalance in gut microbial interactions as a marker of health and disease. Science, 391:890–895, 2026.

[11] A. Eyre-Walker and P. D. Keightley. The distribution of fitness effects of new mutations. Nat Rev Genet, 8:610–618, 2007.

[12] S. Venkataram, H.-Y. Kuo, E. F. Y. Hom, and S. Kryazhimskiy. Mutualism-enhancing mutations dominate early adaptation in a two-species microbial community. Nat Ecol Evol, 7:143–154, 2023.

[13] J. A. Ascensao, K. M. Wetmore, B. H. Good, A. P. Arkin, and O. Hallatschek. Quantifying the local adaptive landscape of a nascent bacterial community. Nat Commun, 14:248, 2023.

[14] J. N. V. Martinson, J. M. Chacón, B. A. Smith, A. R. Villarreal, R. C. Hunter, and W. R. Harcombe. Mutualism reduces the severity of gene disruptions in predictable ways across microbial communities. ISME J, 17:2270–2278, 2023.

[15] E. M. Adamowicz, J. Flynn, R. C. Hunter, and W. R. Harcombe. Cross-feeding modulates antibiotic tolerance in bacterial communities. ISME J, 12:2723–2735, 2018.

[16] B. Pauli, L. Oña, M. Hermann, and C. Kost. Obligate mutualistic cooperation limits evolvability. Nat Commun, 13:337, 2022.

[17] G. R. Lewin, A. Stacy, K. L. Michie, R. J. Lamont, and M. Whiteley. Large-scale identification of pathogen essential genes during coinfection with sympatric and allopatric microbes. Proc Natl Acad Sci USA, 116:19685–19694, 2019.

[18] B. LaSarre, A. M. Deutschbauer, C. E. Love, and J. B. McKinlay. Covert cross-feeding revealed by genome-wide analysis of fitness determinants in a synthetic bacterial mutualism. Appl Environ Microbiol, 86:e00543–20, 2020.

[19] E. C. Pierce, M. Morin, J. C. Little, R. B. Liu, J. Tannous, N. P. Keller, K. Pogliano, B. E. Wolfe, L. M. Sanchez, and R. J. Dutton. Bacterial–fungal interactions revealed by genome-wide analysis of bacterial mutant fitness. Nat Microbiol, 6:87–102, 2021.

[20] J. E. Schreier, C. B. Smith, T. R. Ioerger, and M. A. Moran. A mutant fitness assay identifies bacterial interactions in a model ocean hot spot. Proc Natl Acad Sci USA, 120:e2217200120, 2023.

[21] K. L. Hentchel L. M. Reyes Ruiz, P. D. Curtis, A. Fiebig, M. L. Coleman, and S. Crosson. Genome-scale fitness profile of Caulobacter crescentus grown in natural freshwater. ISME J, 13:523–536, 2019.

[22] W. Shou, S. Ram, and J. M. G. Vilar. Synthetic cooperation in engineered yeast populations. Proc Natl Acad Sci USA, 104:1877–1882, 2007.

[23] M. T. Mee, J. J. Collins, G. M. Church, and H. H. Wang. Syntrophic exchange in synthetic microbial communities. Proc Natl Acad Sci USA, 111:E2149–E2156, 2014.

[24] S. Pande, H. Merker, K. Bohl, M. Reichelt, S. Schuster, L. F. de Figueiredo, C. Kaleta, and C. Kost. Fitness and stability of obligate cross-feeding interactions that emerge upon gene loss in bacteria. ISME J, 8:953–962, 2014.

[25] A. Dal Co, S. van Vliet, D. J. Kiviet, S. Schlegel, and M. Ackermann. Short-range interactions govern the dynamics and functions of microbial communities. Nat Ecol Evol, 4:366–375, 2020.

[26] G. Micali, A. M. Hockenberry, A. Dal Co, and M. Ackermann. Minorities drive growth resumption in crossfeeding microbial communities. Proc Natl Acad Sci USA, 120:e2301398120, 2023.

[27] K. M. Wetmore, M. N. Price, R. J. Waters, J. S. Lamson, J. He, C. A. Hoover, M. J. Blow, J. Bristow, G. Butland, A. P. Arkin, and A. Deutschbauer. Rapid quantification of mutant fitness in diverse bacteria by sequencing randomly bar-coded transposons. mBio, 6:e00306–15, 2015.

[28] B. D. Nguyen, M. Cuenca V., J. Hartl, E. Gül, R. Bauer, S. Meile, J. Rüthi, C. Margot, L. Heeb, F. Besser, P. P. Escriva, C. Fetz, M. Furter, L. Laganenka, P. Keller, L. Fuchs, M. Christen, S. Porwollik, M. McClelland, J. A. Vorholt, U. Sauer, S. Sunagawa, B. Christen, and W.-D. Hardt. Import of aspartate and malate by DcuABC drives H2/fumarate respiration to promote initial Salmonella gut-lumen colonization in mice. Cell Host Microbe, 27:922–936, 2020.

[29] J. W. Fink, D. G. Sant, and M. Manhart. A theoretical framework for how ecological interactions between microbes affect mutant fitness. bioRxiv preprint, doi:TBA, 2026.

[30] J. A. Ascensao, K. D. Abedi, A. N. Prasad, and O. Hallatschek. Frequency-dependent fitness effects are ubiquitous. bioRxiv preprint, doi:10.1101/2025.08.18.670924, 2026.

[31] Ignacio J. Melero-Jiménez, Yael Sorokin, Ami Merlin, Jiawei Li, Alejandro Couce, and Jonathan Friedman. Mutualism breakdown underpins evolutionary rescue in an obligate cross-feeding bacterial consortium. Nature Communications, 16(1):3482, 2025.

[32] Grant Kinsler, Kerry Geiler-Samerotte, and Dmitri A Petrov. Fitness variation across subtle environmental perturbations reveals local modularity and global pleiotropy of adaptation. eLife, 9:e61271, dec 2020.

[33] Encarnación Medina-Carmona, Luis I Gutierrez-Rus, Fadia Manssour-Triedo, Matilda S Newton, Gloria Gamiz-Arco, Antonio J Mota, Pablo Reiné, Juan Manuel Cuerva, Mariano Ortega-Muñoz, Eduardo Andrés-Leon, Jose Luis Ortega-Roldan, Burckhard Seelig, Beatriz Ibarra-Molero, and Jose M Sanchez-Ruiz. Cell survival enabled by leakage of a labile metabolic intermediate. Molecular Biology and Evolution, 40(3):msad032, 03 2023.

[34] Jan Dolinšek, Josep Ramoneda, and David R Johnson. Initial community composition determines the longterm dynamics of a microbial cross-feeding interaction by modulating niche availability. ISME Communications, 2(1):77, 12 2022.

[35] D. Lawrence, F. Fiegna, V. Behrends, J. G. Bundy, A. B. Phillimore, T. Bell, and T. G. Barraclough. Species interactions alter evolutionary responses to a novel environment. PLoS Biol, 10:e1001330, 2012.

[36] E. M. Adamowicz, M. Muza, J. M. Chacón, and W. R. Harcombe. Cross-feeding modulates the rate and mechanism of antibiotic resistance evolution in a model microbial community of Escherichia coli and Salmonella enterica. PLoS Pathog, 16:e1008700, 2020.

[37] Guillaume Martin and Thomas Lenormand. The fitness effect of mutations across environments: A survey in light of fitness landscape models. Evolution, 60(12):2413–2427, 12 2006.

[38] Tim Connallon and Andrew G. Clark. The distribution of fitness effects in an uncertain world. Evolution, 69(6):1610–1618, 06 2015.

[39] J. Jeffrey Morris, Richard E. Lenski, and Erik R. Zinser. The black queen hypothesis: Evolution of dependencies through adaptive gene loss. mBio, 3(2):10.1128/mbio.00036–12, 2012.

[40] Glen D’Souza, Silvio Waschina, Samay Pande, Katrin Bohl, Christoph Kaleta, and Christian Kost. Less is more: Selective advantages can explain the prevalent loss of biosynthetic genes in bacteria. Evolution, 68(9):2559–2570, 09 2014.

[41] Glen D’Souza and Christian Kost. Experimental evolution of metabolic dependency in bacteria. PLOS Genetics, 12(11):1–27, 11 2016.

[42] L. Zhao, L. Zhimin, S. F. Levy, S. Wu, and B. Berger. Bartender: A fast and accurate clustering algorithm to count barcode reads. Bioinformatics, 34:739–747, 2018.

[43] B. Langmead and S. Salzberg. Fast gapped-read alignment with Bowtie 2. Nature Methods, 9:357–359, 2012.

[44] A. Couce, A. Limdi, M. Magnan, S. V. Owen, C. M. Herren, R. E. Lenski, O. Tenaillon, and M. Baym. Changing fitness effects of mutations through long-term bacterial evolution. Science, 383:eadd1417, 2024.

[45] J. W. Fink and M. Manhart. Quantifying microbial fitness in high-throughput experiments. eLife, 13:RP102635, 2024. doi: 10.7554/eLife.102635.1.

