## Supplementary Information for "Amino acid cross-feeding in *E. coli* globally and idiosyncratically alters mutant fitness"

(Dated: May 27, 2026)

* Equal contribution

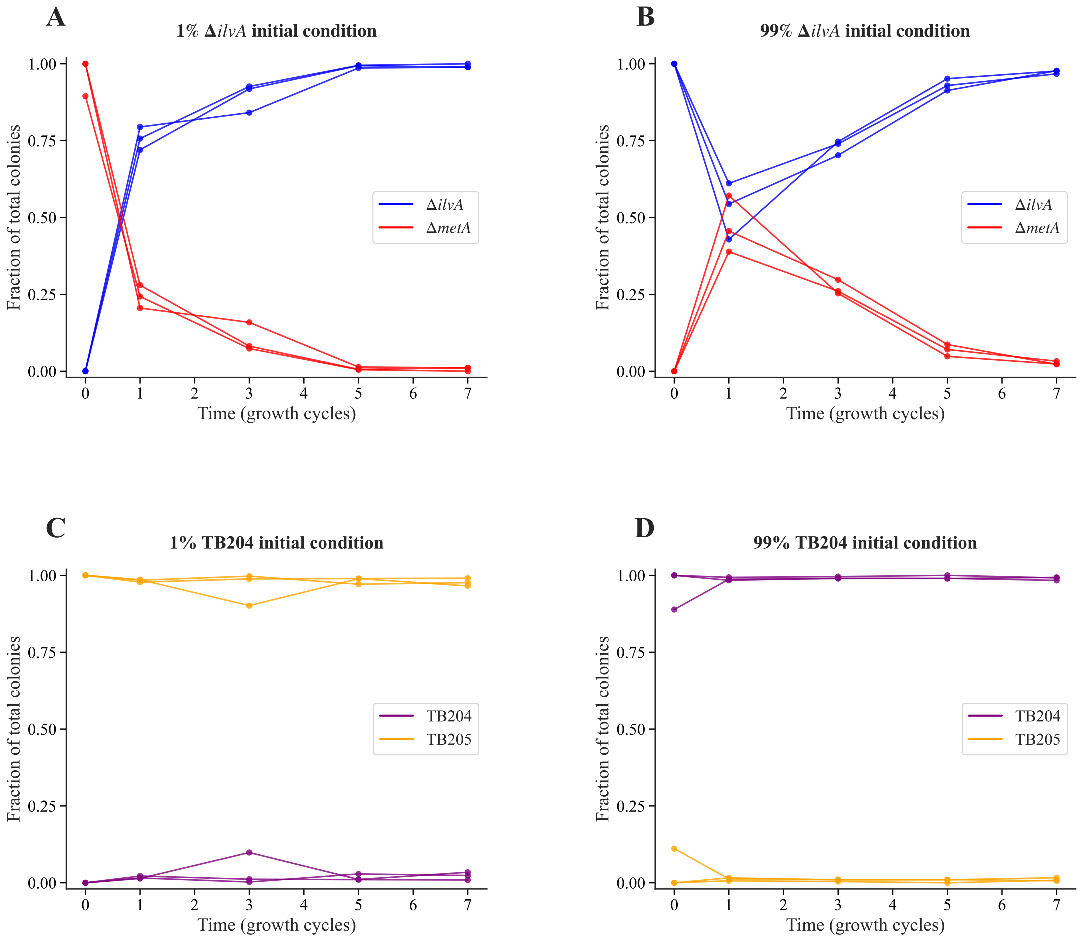


FIG. S1. **Ecological dynamics of the auxotroph cross-feeding community as strain fractions.** Dynamics of $\Delta ilvA$ and $\Delta metA$ across serial transfers starting from (A) 1% $\Delta ilvA$ or (B) 99% $\Delta ilvA$ (Methods). Both converge to a similar composition of 97–99% $\Delta ilvA$. (C, D) Same as (A, B) but for TB204 and TB205, the prototrophic ancestors of $\Delta ilvA$ and $\Delta metA$, as a control to test their neutrality with respect to each other. Figure S2 shows the cell density (CFUs/mL) data from which we derive these strain fractions.

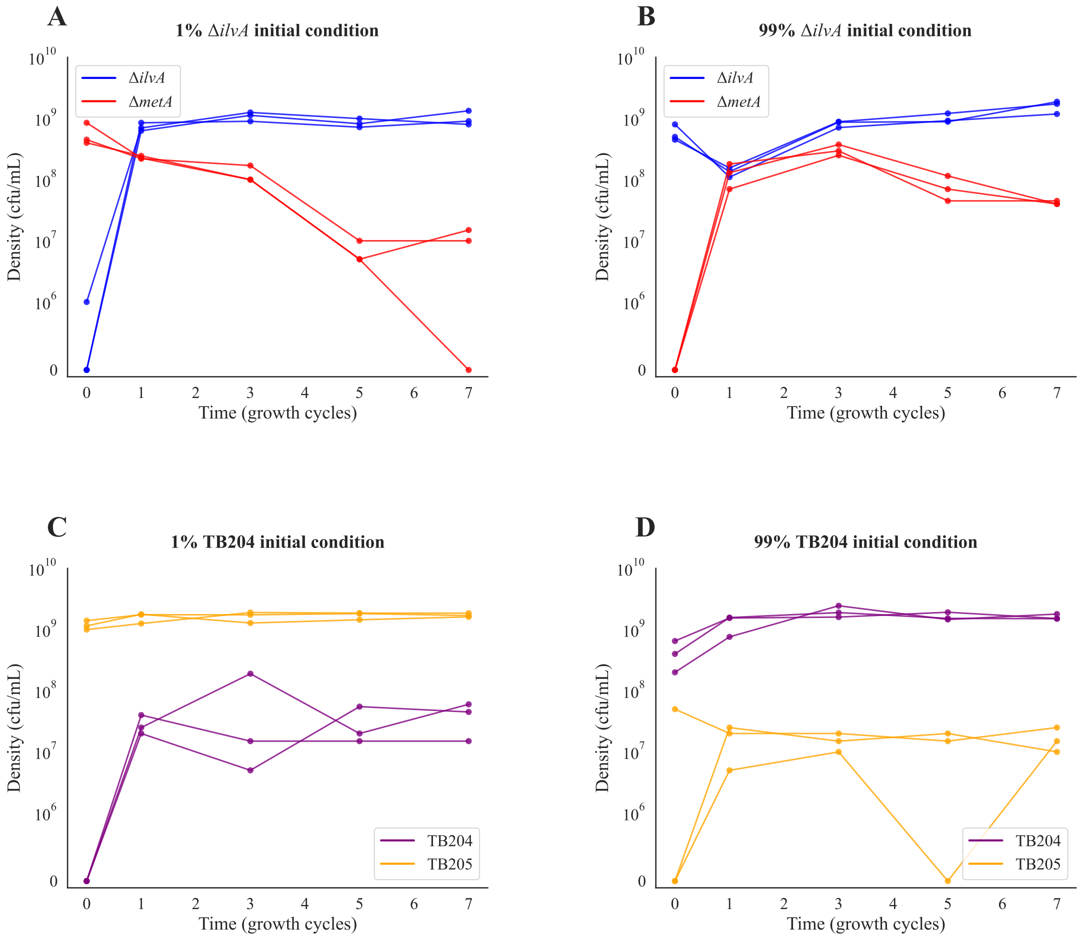


FIG. S2. **Ecological dynamics of the auxotroph cross-feeding community as cell densities.** Same as Fig. S1 but showing cell densities (CFUs/mL) instead of strain fractions (Methods).

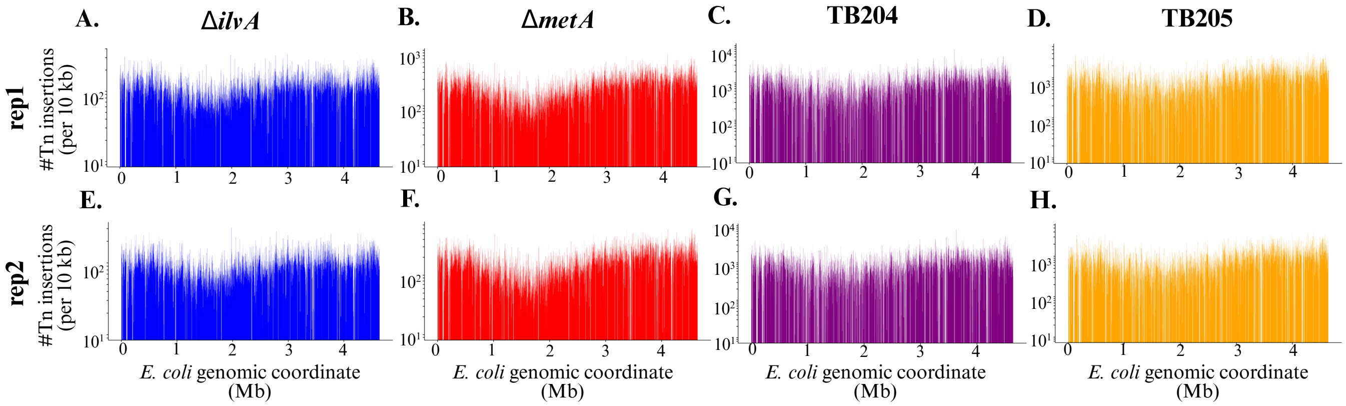


FIG. S3. **Distribution of transposon insertions across genome for each library.** Histograms showing the distribution of transposon insertions (measured as number of transposon insertions per 10 kb) across the *E. coli* genome for $\Delta ilvA$, $\Delta metA$, TB204 (prototrophic ancestor of $\Delta ilvA$ with GFP), and TB205 (prototrophic ancestor of $\Delta metA$ with mCherry), from two replicates of TnSeq (Methods).

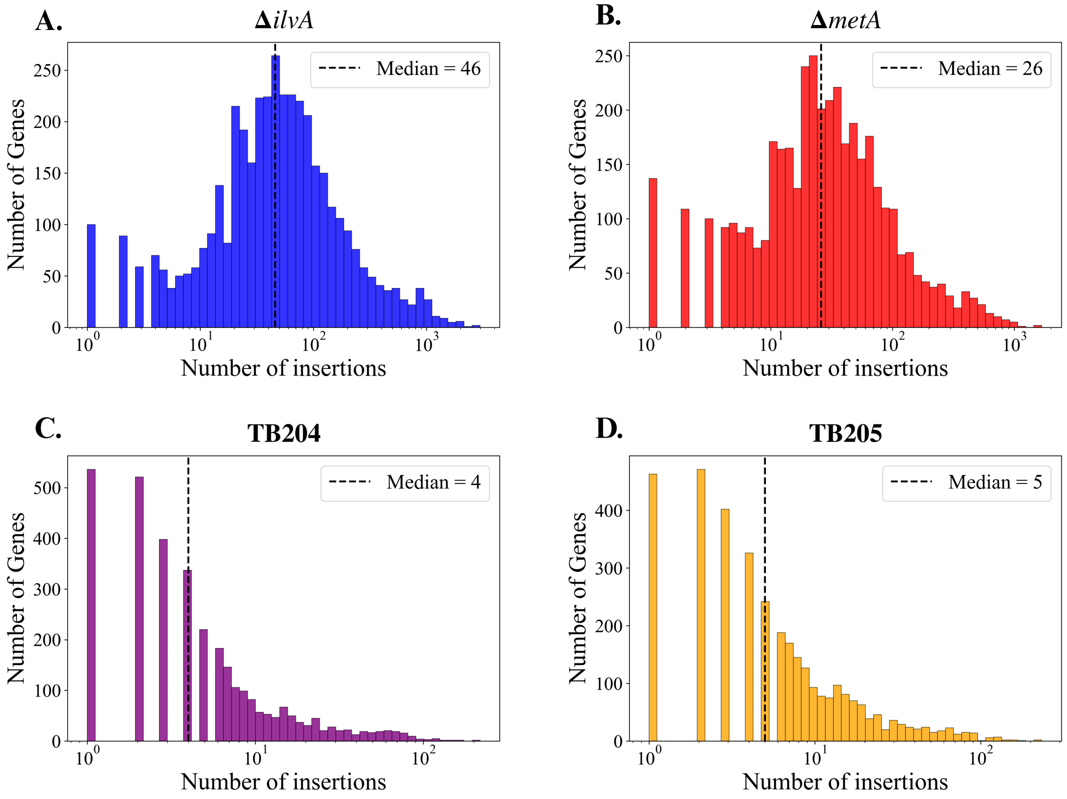


FIG. S4. **Distribution of transposon insertions per gene for each library.** Histograms showing the number of transposon insertions per gene for (A) Δ*ilvA*, (B) Δ*metA*, (C) TB204 (prototrophic ancestor of Δ*ilvA* with GFP), and (D) TB205 (prototrophic ancestor of Δ*metA* with mCherry). The vertical dashed line marks the median in each panel.

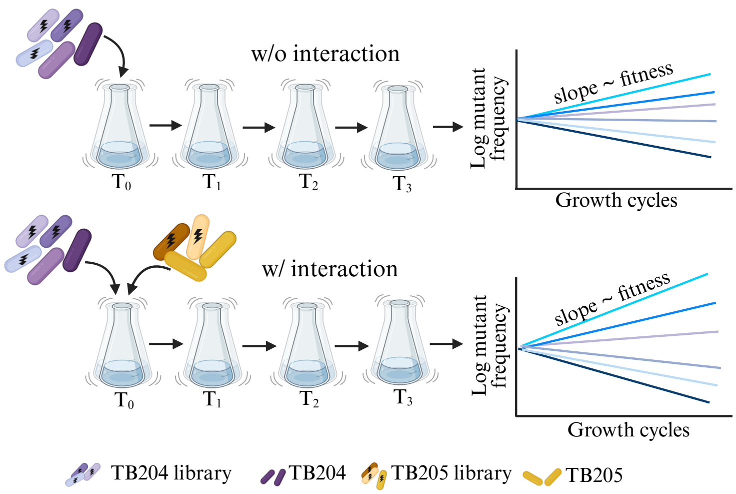


FIG. S5. **Schematic of mutant selection experiments with prototrophs.** Same as main text Fig. 3 but with the prototroph strains.

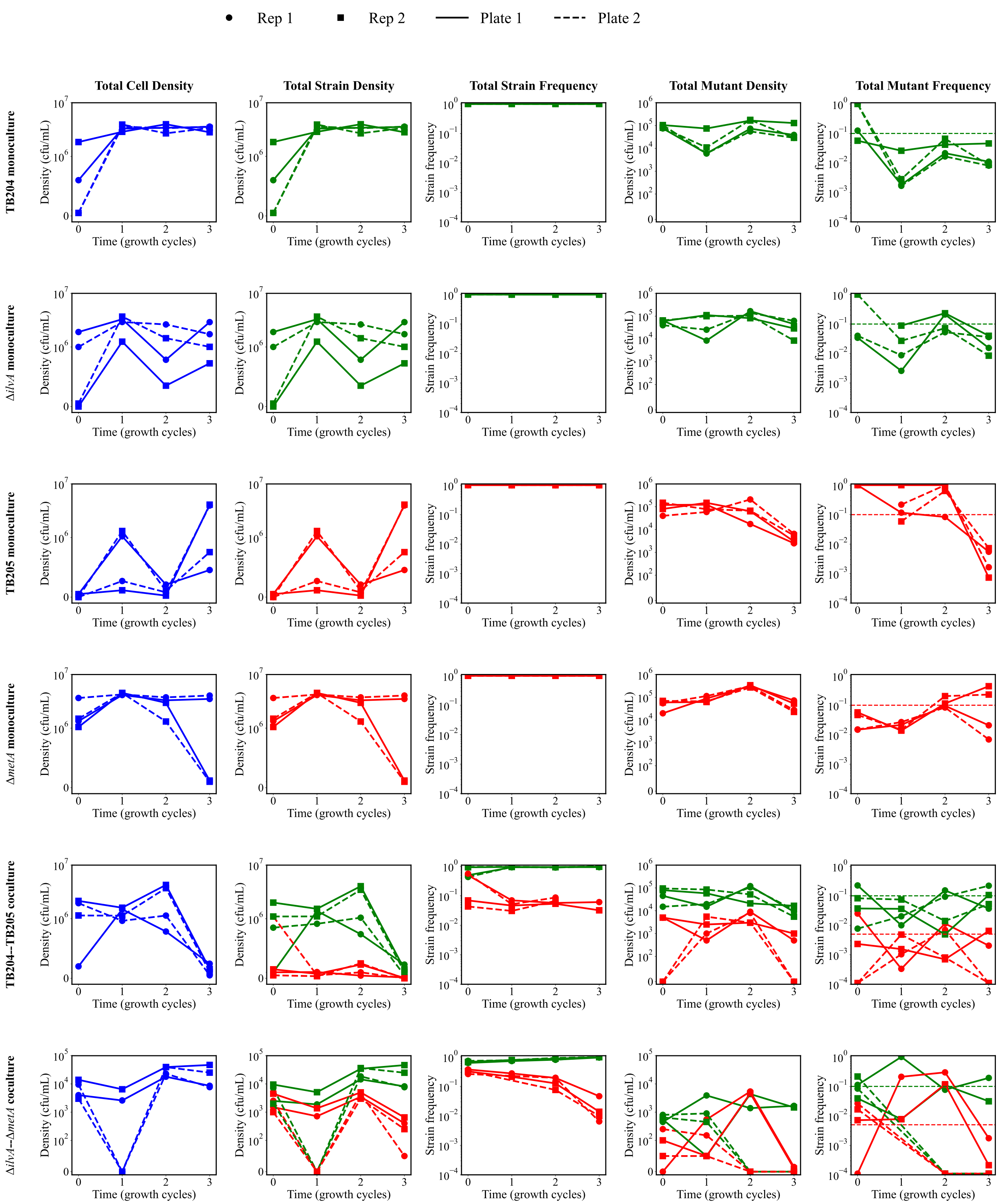


FIG. S6. **Population dynamics of strains and mutant libraries across mutant selection experiments.** Each row corresponds to a mutant selection experiment (main text Fig. 3 and Fig. S5, Methods). The columns report the dynamics of those populations over time as measured by colony counting (Methods). The circle and square points refer to the two biological replicate experiments, and the solid and dashed lines refer to replicate plate measurements for each. First column is total cell density (CFUs/mL) of all strains in the culture. Second column is cell density of the GFP- ($\Delta ilvA$ or TB204) and the mCherry-containing ($\Delta metA$ or TB205) strains. Third column is the same data as the second column, but normalized as fractions. Fourth column is the cell density of the GFP- ($\Delta ilvA$ or TB204) and the mCherry-containing ($\Delta metA$ or TB205) mutant libraries as wholes. Fifth column is the fraction of the GFP- ($\Delta ilvA$ or TB204) and the mCherry-containing ($\Delta metA$ or TB205) mutant libraries relative to all GFP- or mCherry-containing cells (e.g., data in fourth column normalized by the data in the second column).

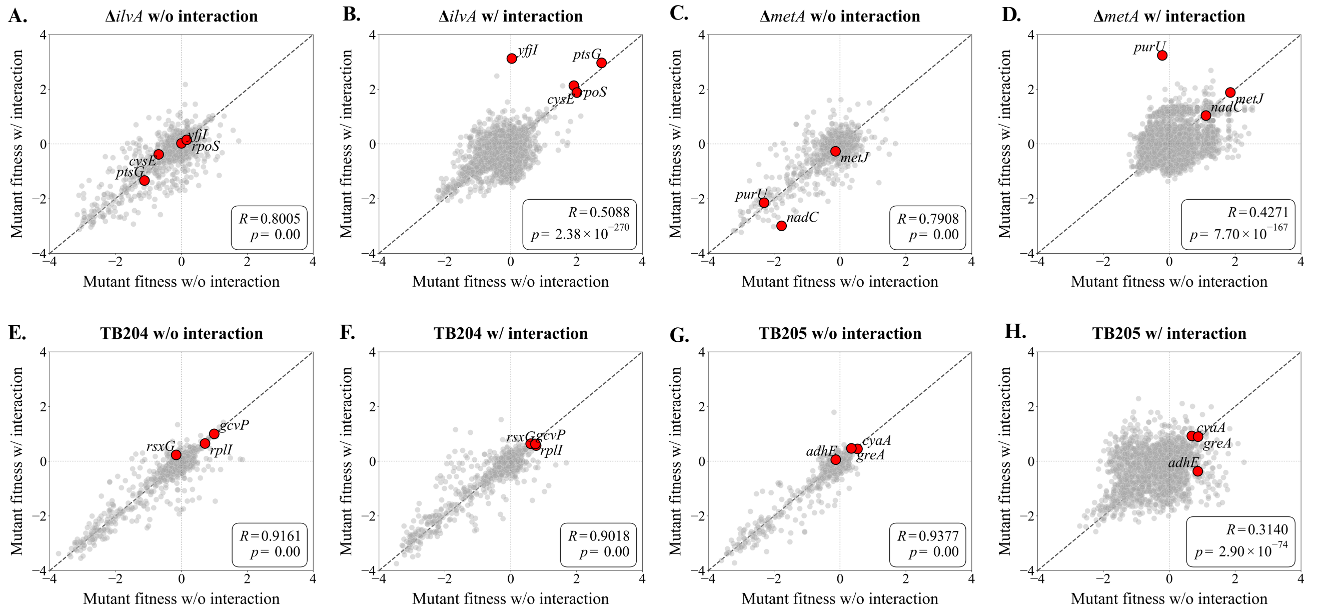


FIG. S7. **Correlation of fitness measurements across biological replicates.** Each panel shows a scatter plot of fitness from two biological replicates of the labeled selection experiments for all mutants (individual insertions in a single gene aggregated together; Methods). The legends show the Pearson correlation coefficients $R$ and their associated $p$-values; the black diagonal line is the identity. Red points mark genes with large beneficial effects in the cocultures that we investigate in main text Figs. 5 and 6, as well as Figs. S10, S11, and S12.

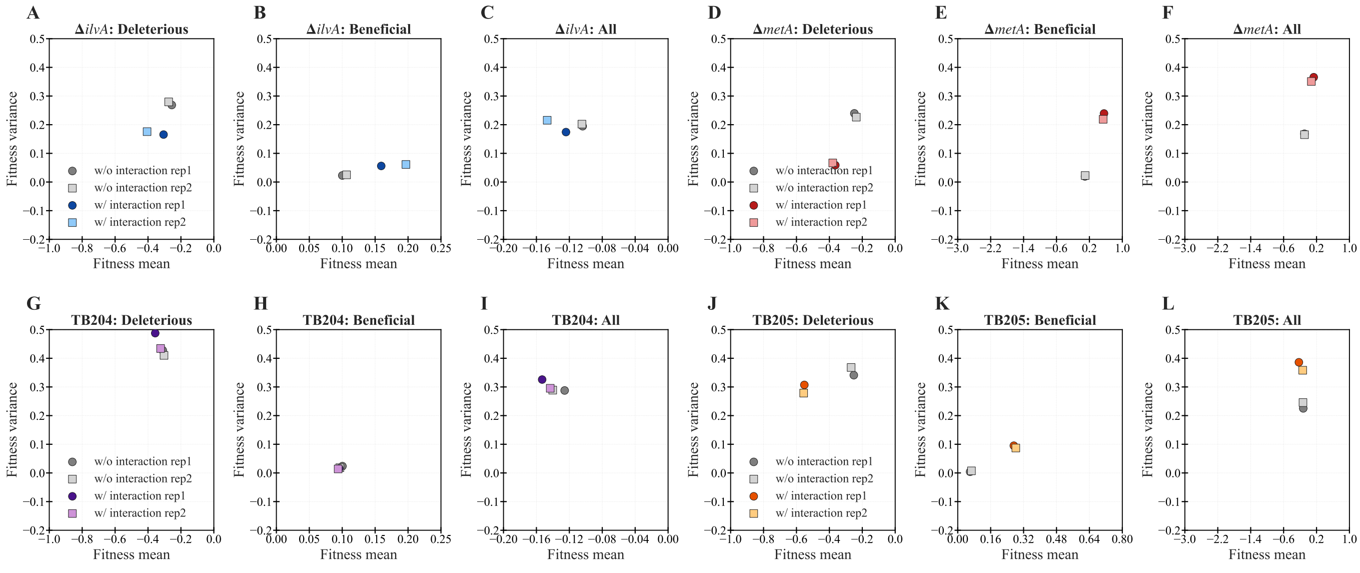


FIG. S8. **Variation in DFE mean and variance across replicates and culture conditions.** Each panel corresponds to a mutant library and shows the mean and variance of the DFE for the library across the two replicate selection experiments and the culture conditions with and without the interaction. The sets of three panels for the same library break down these statistics for deleterious mutants only, beneficial mutants only, and for all mutants.

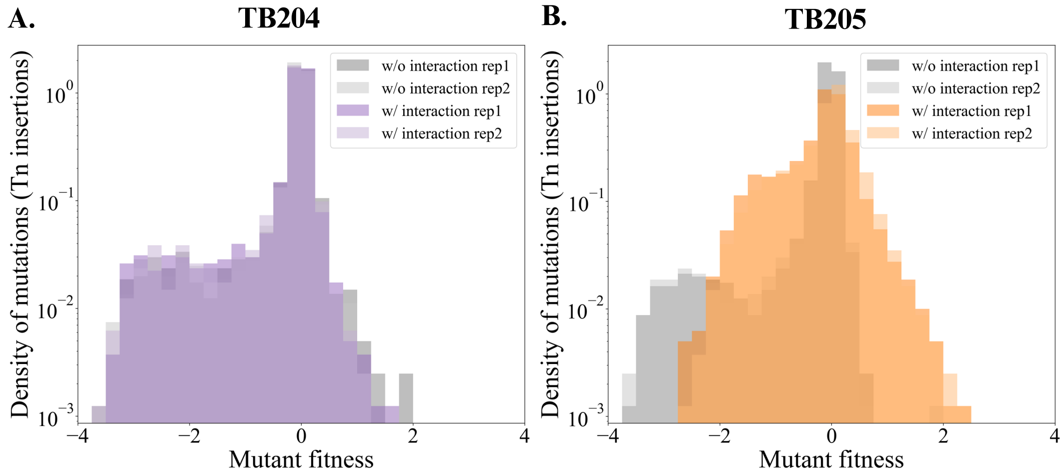


FIG. S9. **Testing global effects on DFEs for prototroph strains.** Same as main text Fig. 4 but for the prototroph strains TB204-GFP (ancestor of $\Delta ilvA$) and TB205-mCherry (ancestor of $\Delta metA$).

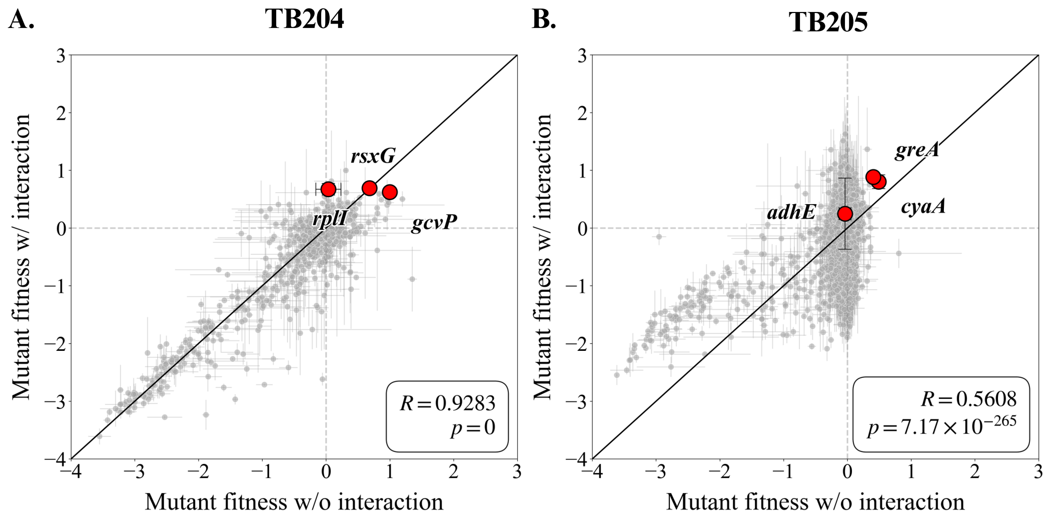


FIG. S10. **Testing idiosyncratic effects on DFEs for prototroph stains.** Same as main text Fig. 5 but for the prototroph strains TB204-GFP (ancestor of $\Delta ilvA$) and TB205-mCherry (ancestor of $\Delta metA$).

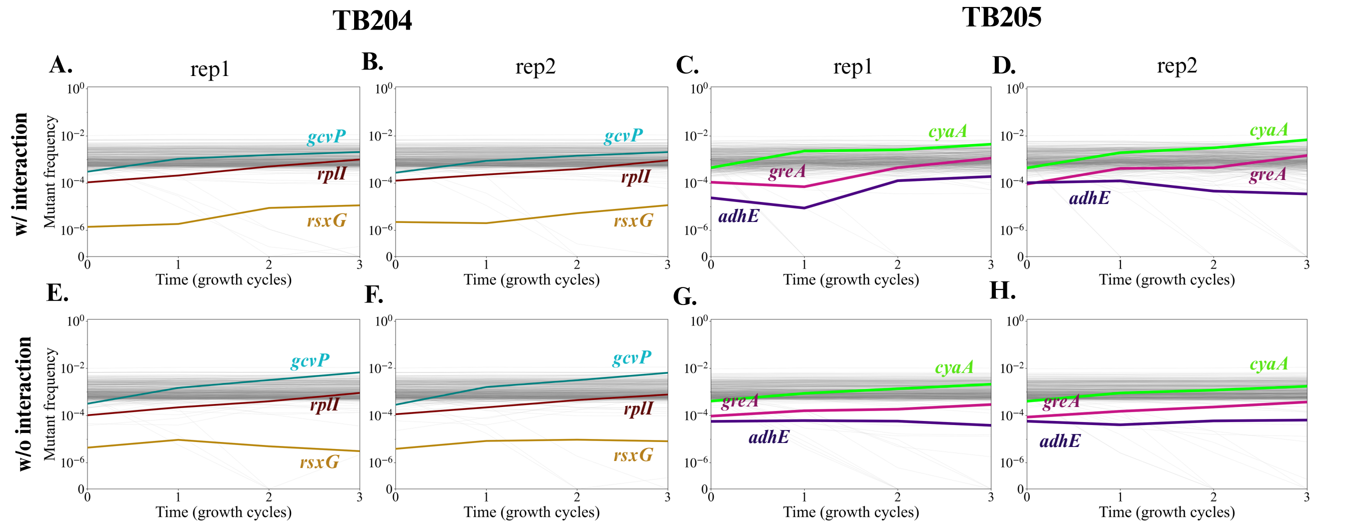


FIG. S11. **Dynamics of individual mutants from prototroph strains.** Same as main text Fig. 6 but for the prototroph strains TB204-GFP (ancestor of $\Delta ilvA$) and TB205-mCherry (ancestor of $\Delta metA$).

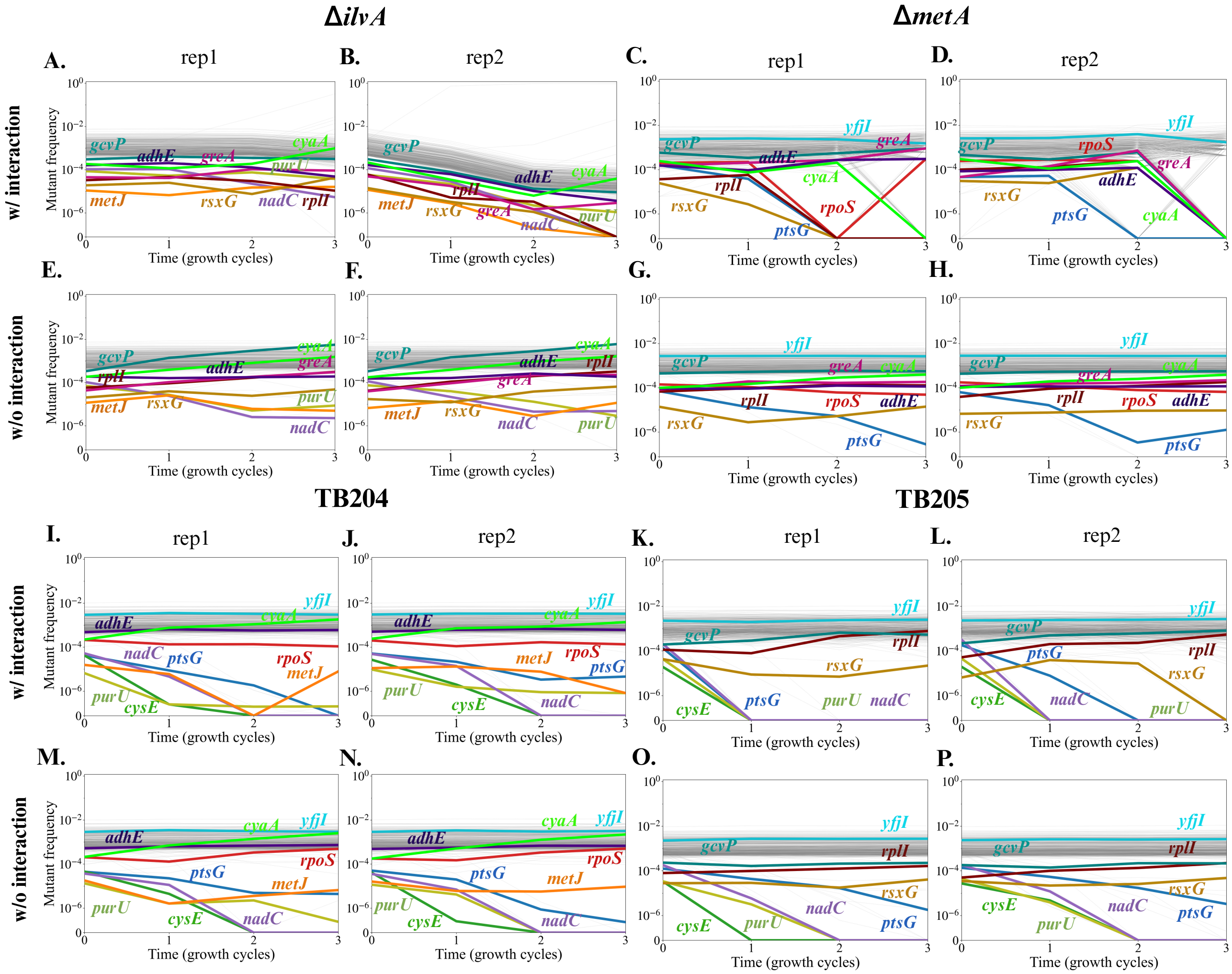


FIG. S12. **Dynamics of top mutants from cocultures in other genetic backgrounds.** For the most beneficial mutants from the coculture conditions highlighted in main text Fig. 6 and Fig. S11, we plot the dynamics of insertions in those same genes but on the backgrounds of the three other strains in our set of four mutant libraries. Format is otherwise the same as main text Fig. 6 and Fig. S11.

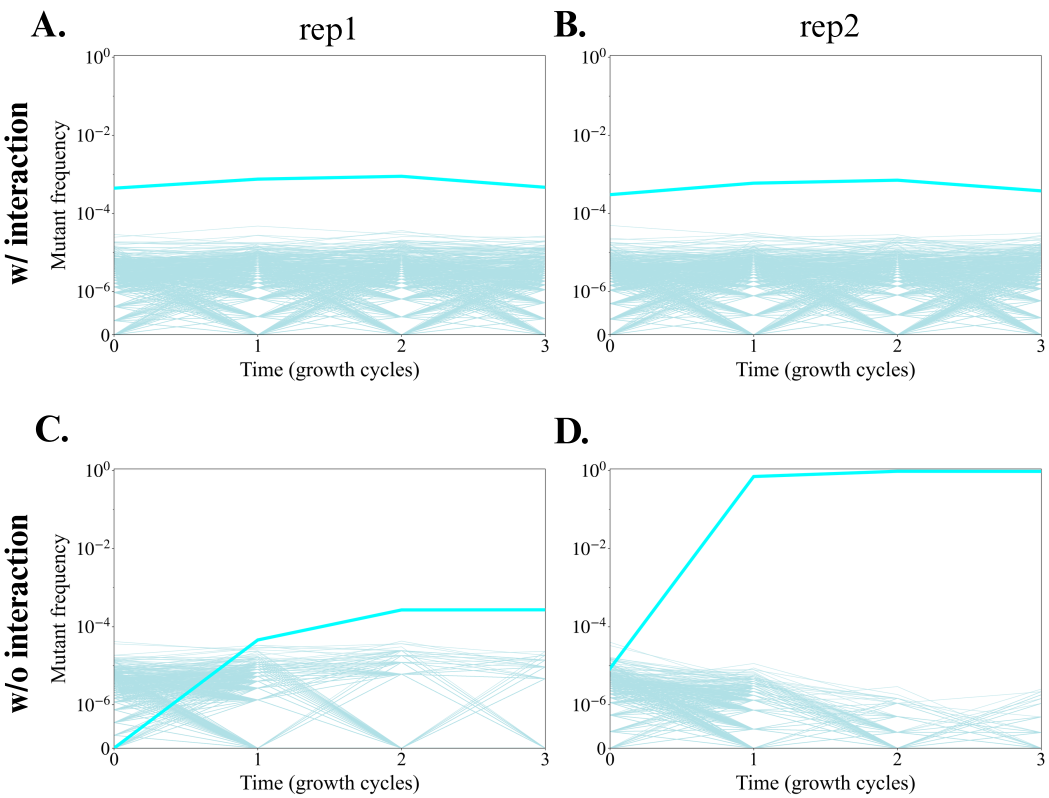


FIG. S13. **Dynamics of all insertions in *yfjI* in the Δ*ilvA* mutant library.** Each panel shows trajectories of all 789 insertions in *yfjI* in both biological replicates of the mutant selection experiment and the two culture conditions. The thick line is the single insertion that rises to high frequency in replicate 2 of the cross-feeding condition.

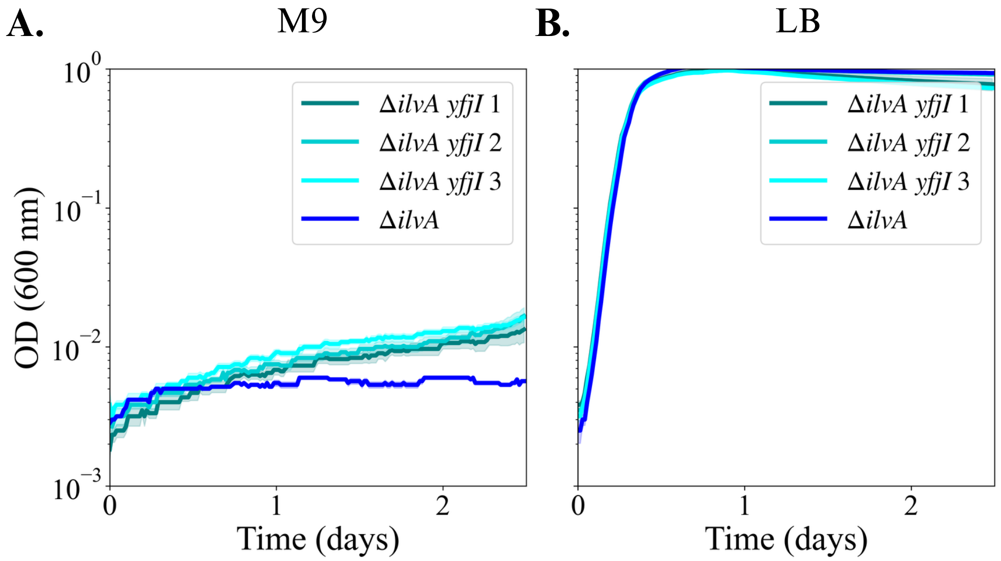


FIG. S14. **Negative and positive controls of growth curves for *yfjI* insertion strain and its Δ*ilvA* ancestor.** Same as main text Fig. 7A–C but showing growth curves in (A) minimal media without any supplemented amino acids and (B) rich media (Methods).

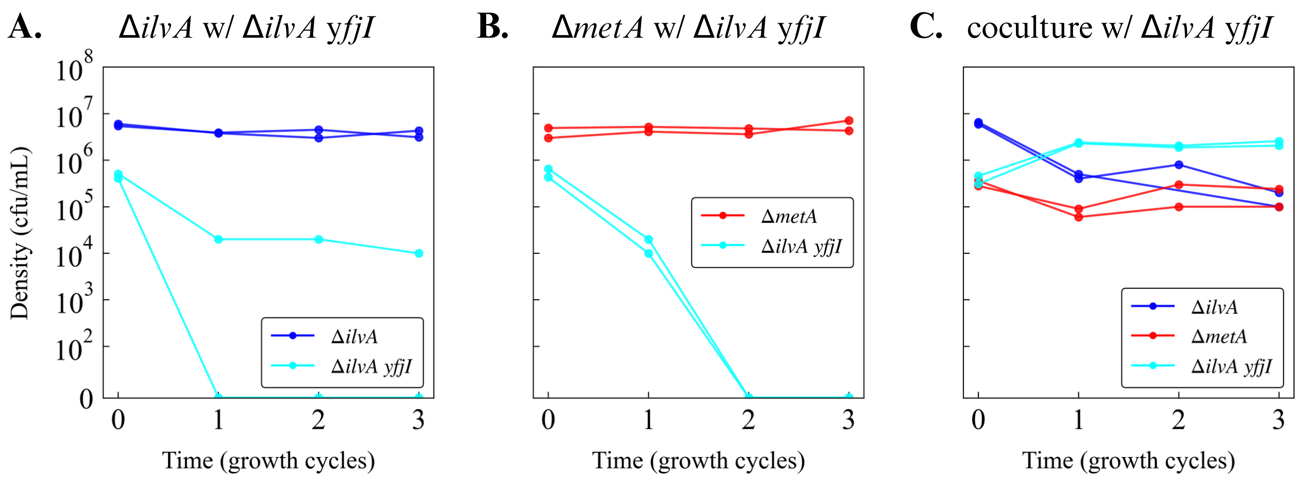


FIG. S15. **Dynamics of additional selection experiments with isolates of the *yfjI* insertion strain.** Same as main text Fig. 7D–F but showing cell density (CFUs/mL) from which we derive the strain fractions in main text Fig. 7D–F.

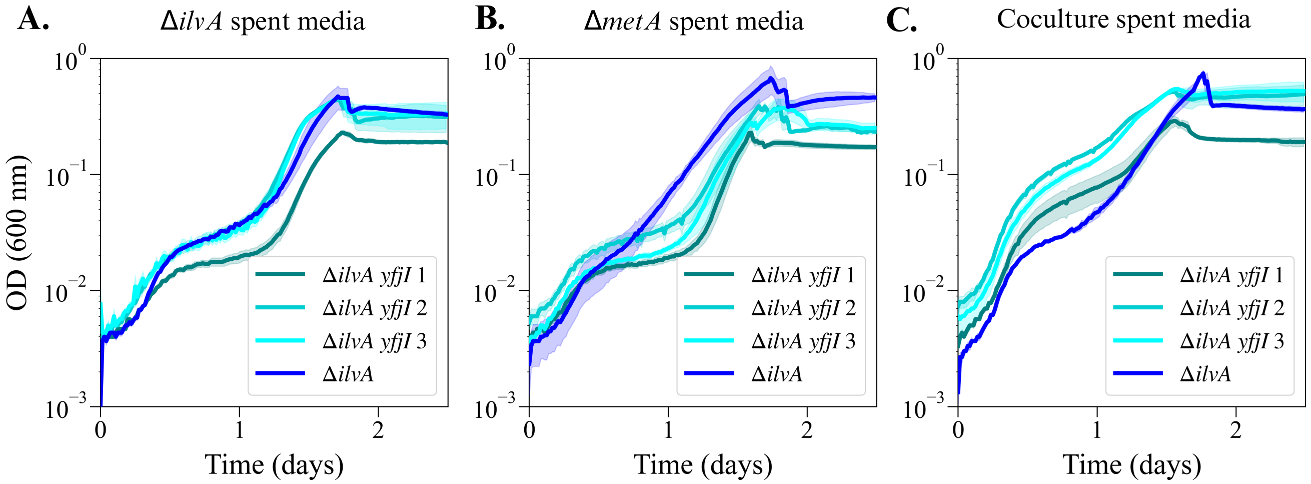


FIG. S16. **Growth of *yfjI* insertion strains in different spent media.** Growth curves of three isolates of the *yfjI* insertion strain and the Δ*ilvA* ancestor in minimal media supplemented with spent media from (A) Δ*ilvA* monoculture, (B) Δ*metA* monoculture, and (C) Δ*ilvA* and Δ*metA* coculture (Methods). Dark line represents the mean and shading the standard error from three technical replicates (same inoculum).

| Name | Description | Antibiotic marker |
|---|---|---|
| Δ*ilvA* | *E. coli* K-12 MG1655 - Δ*ilvA*-GFP | no |
| Δ*metA* | *E. coli* K-12 MG1655 - Δ*metA*-mCherry | no |
| TB204 | *E. coli* K-12 MG1655 - TB204-GFP | no |
| TB205 | *E. coli* K-12 MG1655 - TB205-mCherry | no |
| Δ*ilvA yfjI* mutant | *E. coli* K-12 MG1655 - Δ*ilvA*-GFP with *yfjI* Tn5 insertion | Kanamycin |

TABLE S1. **Strains used for the experiments.**

| | Selection experiment culture condition | | | | | |
|---|---|---|---|---|---|---|
| **Component** | **Δ*ilvA* monoculture** | **Δ*metA* monoculture** | **TB204 monoculture** | **TB205 monoculture** | **Δ*ilvA* - Δ*metA* coculture** | **TB204 and TB205 coculture** |
| 1X M9 salts | 147.51 mL | 147.51 mL | 147.51 mL | 147.51 mL | 147.81 mL | 147.81 mL |
| 1M MgSO4 | 300 µL | 300 µL | 300 µL | 300 µL | 300 µL | 300 µL |
| 1M CaCl2 | 15 µL | 15 µL | 15 µL | 15 µL | 15 µL | 15 µL |
| 200 g/L Glucose | 375 µL | 375 µL | 375 µL | 375 µL | 375 µL | 375 µL |
| 50 g/L amino acid | 300 µL Ile | 300 µL Met | 300 µL Ile | 300 µL Met | - | - |

TABLE S2. **Composition of M9 media used for mutant selection experiments.**

| Library | Comparison Type | Sample A | Sample B | KS Statistic | p-value |
|---|---|---|---|---|---|
| Δ*ilvA* mutant library | Replicate Comparison (Mono) | monoR1_fitness | monoR2_fitness | 0.0306 | 4.2449e-02 |
| Δ*ilvA* mutant library | Replicate Comparison (Co) | coR1_fitness | coR2_fitness | 0.0803 | 5.5440e-12 |
| Δ*ilvA* mutant library | Treatment Comparison (Rep 1) | monoR1_fitness | coR1_fitness | 0.1344 | 7.2828e-33 |
| Δ*ilvA* mutant library | Treatment Comparison (Rep 2) | monoR2_fitness | coR2_fitness | 0.1815 | 1.0387e-59 |
| Δ*metA* mutant library | Replicate Comparison (Mono) | monoR1_fitness | monoR2_fitness | 0.0218 | 3.3374e-01 |
| Δ*metA* mutant library | Replicate Comparison (Co) | coR1_fitness | coR2_fitness | 0.0462 | 6.4254e-04 |
| Δ*metA* mutant library | Treatment Comparison (Rep 1) | monoR1_fitness | coR1_fitness | 0.3434 | 3.1469e-197 |
| Δ*metA* mutant library | Treatment Comparison (Rep 2) | monoR2_fitness | coR2_fitness | 0.3102 | 2.1504e-160 |
| TB204 mutant library | Replicate Comparison (Mono) | monoR1_fitness | monoR2_fitness | 0.05 | 6.3686e-04 |
| TB204 mutant library | Replicate Comparison (Co) | coR1_fitness | coR2_fitness | 0.0255 | 2.4737e-01 |
| TB204 mutant library | Treatment Comparison (Rep 1) | monoR1_fitness | coR1_fitness | 0.0295 | 1.2126e-01 |
| TB204 mutant library | Treatment Comparison (Rep 2) | monoR2_fitness | coR2_fitness | 0.0366 | 2.6480e-02 |
| TB205 mutant library | Replicate Comparison (Mono) | monoR1_fitness | monoR2_fitness | 0.0234 | 3.4386e-01 |
| TB205 mutant library | Replicate Comparison (Co) | coR1_fitness | coR2_fitness | 0.1186 | 4.8405e-20 |
| TB205 mutant library | Treatment Comparison (Rep 1) | monoR1_fitness | coR1_fitness | 0.2656 | 9.2295e-100 |
| TB205 mutant library | Treatment Comparison (Rep 2) | monoR2_fitness | coR2_fitness | 0.2768 | 1.8531e-108 |

TABLE S3. **Statistics of Kolmogorov-Smirnov tests between pairs of DFEs.**

| **Position** | **Mutation** | **Δ*ilvA* rep 1** | **Δ*ilvA* rep 2** | **Δ*ilvA* rep 3** | **Δ*ilvA* *yfjI* rep1** | **Δ*ilvA* *yfjI* rep2** | **Δ*ilvA* *yfjI* rep3** | **Annotation** | **Gene** | **Description** |
|---|---|---|---|---|---|---|---|---|---|---|
| 4,127 | G→C | | | | 100% | 100% | 100% | A132P (GCG→CCG) | *thrC* → | threonine synthase |
| 1,196,220 | C→T | 100% | | 100% | 100% | 100% | 100% | H366H (CAC→CAT) | *icd* → | isocitrate dehydrogenase |
| 1,196,226 | T→C | | | | | | 100% | G368G (GGT→GGC) | *icd* → | isocitrate dehydrogenase |
| 1,196,232 | C→T | | | | | | 100% | T370T (ACC→ACT) | *icd* → | isocitrate dehydrogenase |
| 1,812,021 | G→A | 100% | 100% | 100% | 100% | 100% | 100% | G363D (GGC→GAC) | *tcyP* → | cystine/sulfocysteine:cation symporter |
| 1,978,503 | Δ776 bp | 100% | | 100% | | | | | *insB5–insA5* | *insB5*, *insA5* |
| 2,173,363 | Δ2 bp | 100% | 100% | 100% | 100% | 100% | 100% | intergenic (-490/+918) | *gatD* ← / ← *gatB* | galactitol-1-phosphate 5-dehydrogenase/galactitol-specific PTS enzyme IIB component |
| 2,760,199 | Δ1,452 bp | 100% | 100% | 100% | | | | coding (1215-2666/2862 nt) | *yfjI* → | DUF3987 domain-containing protein YfjI |
| 2,761,332 | +G | | | | 100% | 100% | 100% | coding (2348/2862 nt) | *yfjI* → | DUF3987 domain-containing protein YfjI |
| 2,761,342 | Δ1 bp | | | | 100% | 100% | 100% | coding (2358/2862 nt) | *yfjI* → | DUF3987 domain-containing protein YfjI |
| 2,761,650 | +GCTTTATTG | | | | 100% | 100% | 100% | coding (2666/2862 nt) | *yfjI* → | DUF3987 domain-containing protein YfjI |
| 3,561,907 | +G | 100% | 100% | 100% | 100% | 100% | 100% | intergenic (+589/+167) | *rtcR* → / ← *glpG* | DNA-binding transcriptional activator RtcR/rhomboid protease GlpG |
| 4,297,833 | +GC | 100% | 100% | 100% | 100% | 100% | 100% | intergenic (+587/+55) | *gltP* → / ← *yjcO* | glutamate/aspartate : H(+) symporter GltP/Sel1 repeat-containing protein YjcO |

TABLE S4. **Genomic variants in *yfjI* insertion strain isolates and Δ*ilvA* isolates.**
